# Dynamic Functional Connectivity between Order and Randomness and its Evolution across the Human Adult Lifespan

**DOI:** 10.1101/107243

**Authors:** Demian Battaglia, Thomas Boudou, Enrique C. A. Hansen, Diego Lombardo, Sabrina Chettouf, Andreas Daffertshofer, Anthony R. McIntosh, Joelle Zimmermann, Petra Ritter, Viktor Jirsa

**Author notes:** Shared last authorship (Petra Ritter, Viktor Jirsa). Correspondence to: Demian Battaglia. Other e-mail addresses: Thomas Boudou, Enrique C. A. Hansen, Diego Lombardo, Sabrina Chettouf, Andreas Daffertshofer, A. Randal McIntosh, Joelle Zimmermann, Petra Ritter, Viktor Jirsa.

## Abstract

Functional Connectivity (FC) during resting-state or task conditions is not fixed but inherently dynamic. Yet, there is no consensus on whether fluctuations in FC may resemble isolated transitions between discrete FC states rather than continuous changes. This quarrel hampers advancing the study of dynamic FC. This is unfortunate as the structure of fluctuations in FC can certainly provide more information about developmental changes, aging, and progression of pathologies. We merge the two perspectives and consider dynamic FC as an ongoing network reconfiguration, including a stochastic exploration of the space of possible steady FC states. The statistical properties of this random walk deviate both from a purely “order-driven” dynamics, in which the mean FC is preserved, and from a purely “randomness-driven” scenario, in which fluctuations of FC remain uncorrelated over time. Instead, dynamic FC has a complex structure endowed with long-range sequential correlations that give rise to transient slowing and acceleration epochs in the continuous flow of reconfiguration. Our analysis for fMRI data in healthy elderly revealed that dynamic FC tends to slow down and becomes less complex as well as more random with increasing age. These effects appear to be strongly associated with age-related changes in behavioural and cognitive performance.

**Highlights:**

- Dynamic Functional Connectivity (dFC) at rest and during cognitive task performs a “complex” (anomalous) random walk.
- Speed of dFC slows down with aging.
- Resting dFC replaces complexity by randomness with aging.
- Task performance correlates with the speed and complexity of dFC.

## Introduction

Recent studies emphasised the structured temporal variability of resting-state (rs) and task Functional Connectivity (FC; Tagliazucchi et al., 2012; Allen et al., 2012; Liu & Duyn, 2013; Hutchison et al., 2013; Chen et al., 2015; Preti et al., 2017; Gonzalez-Castillo & Bandettini, 2018), whose study is the defining focus of a research direction recently designated as “chronnectomics” (Calhoun et al., 2014). If rs-FC is dynamic, a wealth of information may be lost by averaging over long imaging sessions, and averaged temporal variability might be mistaken as inter-subject variability. Temporal FC variability –which we will refer to as *dynamic FC* (dFC)– may carry an inherent meaning. It has been suggested to manifest ongoing cognition at rest (Gonzalez-Castillo et al., 2015) with an immediate impact on cognitive performance (Bassett et al., 2011; Braun et al., 2015; Shine et al., 2016; Cohen, 2018) and attentional or awareness levels (Kucyi et al., 2017; Cavanna et al., 2018; Lim et al., 2018). It may reflect the sampling of an internal repertoire of alternative dynamical states (Hansen et al., 2015; Golos et al., 2015; Cabral et al., 2017a; Glomb et al., 2017). From a biomarking perspective, pathological conditions, such as Alzheimer’s dementia, schizophrenia and other mental disorders (Jones et al., 2012; Damaraju et al., 2014; Braun et al., 2018), or differences in general attributes like gender (Yaesoubi et al., 2015) or development and aging (Hutchison & Morton, 2015; Qin et al., 2015; Davison et al., 2016; Schlesinger et al., 2016; Chen et al., 2017; Viviano et al., 2017), may alter dFC more than they affect time-averaged FC.

The growth in the number of dFC studies, based on both fMRI and electrophysiological signals, has been paralleled by an increasing number of possible technical approaches to estimate dFC (Preti et al., 2017). A non-exhaustive list ranges from sliding window approaches (Allen et al., 2012), to statistical modelling of signals (Lindquist et al., 2014) and state transitions (Baker et al., 2014; Vidaurre et al., 2016; Cabral et al., 2017b), temporal network approaches (Thompson & Fransson, 2016), or the study of “coactivation patterns” (“CAPs”: Chen et al., 2015; Matsui et al., 2016). However, several concerns have been raised on whether dFC reflects genuine neural network dynamics or rather artefactual fluctuations, linked, e.g., to head motion (Laumann et al., 2016) or signal processing aspects (Leonardi & Van de Ville, 2015). There are also several statistical concerns about whether resting-state FC is really non-stationary (Zaleski et al, 2014; Hindriks et al., 2016) or whether discrete connectivity states exist that might be reliably extracted (Shakil et al., 2016; Liégeois et al., 2017). In fact, while “FC clusters” can always be extracted using ad hoc algorithmic methods, as of yet that is not evident that such clusters correspond to well-defined, distinct attractor states (Zaleski & Breakspear, 2015).

Here, we introduce yet another way to look at dFC, which, we believe, circumvents some of the concerns mentioned above on the difficulty of assessing the actual non-stationarity of FC fluctuations in time. We do not attempt segmenting dFC in a sequence of sharp switching transitions between FC states but instead describe it as a smooth flow across continually morphing connectivity configurations. Conventional analyses of static FC emphasise the spatial structure of FC networks discarding most information about time. We adopt the opposite approach: de-emphasizing space and collapsing FC networks to a “point” in the space of possible FC network realisations. And, we interpret its erratic evolution as a random walk, a stochastic exploration of a high-dimensional space. An iconic example of random walk is the one of a grain of dust floating in the air, made visible through a beam of shining light. By sampling the position of this grain at different moments in time, one may observe that the displacements of the grain over a fixed observation interval are not constant in time. On the contrary, the distances travelled over different time intervals (or, equivalently, the speeds of motion) give rise to a distribution, which will be (close to) a Gaussian with a well-defined peak. Thus, the mean *speed* of the moving particle constitutes a first metric for the quantitative characterization of its random walk. As a second step, one can consider the statistical properties of the fluctuations of random walk speed around this mean. Not all the random walks are identical and exploration paths with different *shapes* (i.e., fractal geometries) might be generated, depending on the degree of correlation between the step lengths travelled at consecutive times (Mandelbrot & Van Ness, 1968; Mandelbrot, 1983). In some random walks –such as the one of a grain of dust, or of a particle suspended in water, originally studied by Robert Brown (1828)– the speed of consecutive exploration steps are uncorrelated (“memoryless”), giving rise to unstructured uniformly space-filling paths. In some other cases –such as the spread of infectious diseases in contemporary times (Brockmann et al., 2006) or foraging for resources in an ecosystem (Viswanathan et al., 1999)–, consecutive random walk steps have correlated speeds (“memory”), giving rise to paths with a characteristic alternation between exploration of localized clusters and longer jumps. Methods like detrended fluctuation analysis (DFA, Peng et al., 1993) serve to characterize the geometry of empirically observed random walk paths, so to estimate their degree of memory and their deviation from Gaussian uncorrelated walks. Such approaches have been successfully applied to detect anomalous scaling properties of fluctuations in neural datasets at different spatial and temporal scales (He, 2014), associated to critical behaviour in both rest and task conditions (Linkenkaer-Hansen et al., 2001; Chialvo, 2010; Palva et al., 2013). Here, with a picture in mind of resting state dFC has a random walk, we will study its typical speed and the shape of exploration paths in FC space.

We extend previous results that indicated how ongoing FC fluctuations implement a complex random walk, endowed with non-trivial statistical properties. The features of this dFC random walk appear to be intermediate between two possible trivial null hypotheses: a first “order” scenario in which stationarity is strictly imposed, and a second “randomness” scenario in which the time ordering of the observed time-resolved FC matrices is shuffled to destroy any long-range sequential correlations. For FC time series dominated by stochastic properties, a direct means of characterizing such scenarios of time-dependent observations is. Using DFA analysis and statistical comparisons with suitable surrogate dFC streams, we find that dFC deviates from both “order” and “randomness”, being thus “complex” (Crutchfield, 2011).

We also investigate how random walk properties of rest and task dFC may be modified over the healthy human adult lifespan. As we age, our brain undergoes characteristic structural and functional changes, with a tendency toward increased structural ‘disconnection’ (Salat, 2011), disruptions in rs-FC (Andrews-Hanna et al., 2007; Betzel et al., 2014) and modified structural-to-functional connectivity inter-relations (Zimmermann et al., 2016). Analogously, changes in dFC have been reported at the level of the temporal stability of FC network modules (Davison et al., 2016; Schlesinger et al., 2016), general or specific network variability (Qin et al., 2015; Chen et al., 2017), “FC state” occupancy (Hutchison & Morton, 2015; Viviano et al., 2017), and complexity of phase synchrony (Nobukawa et al., 2019). We complement these previous findings and show that dFC random walks may occur at an increasingly reduced speed and complexity with age, slowing down and becoming increasingly more “random”. These reductions in dFC speed and complexity correlate with the level of general cognitive and behavioural performance, as probed by both standard clinical assessments of cognitive impairments (Nasreddine et al., 2005) and a simple visuomotor coordination task (Houweling et al., 2008). For a refined analysis of dFC random walk alterations along with more subtle and task-specific cognitive decline, see a related paper by Lombardo et al. (2020). The fact that slowing down and complexity loss in dFC are associated with degraded general performance can be linked to prominent theories of cognitive aging, speculatively establishing alterations of dFC random walk properties as novel imaging correlates of processing speed reduction (Salthouse, 1996; Finkel et al., 2007) and de-differentiation (Baltes, 1980; Sleimen-Malkoun et al., 2014).

## Materials and methods

### Experimental subjects

Overall *N* = 85 healthy adult subjects (*N* = 53 females, *N* = 32 males) were recruited at Charité - Universitätsmedizin Berlin to voluntarily participate in rs-fMRI and DSI scans and, for a subset of them, also in a visuomotor study. The first subset of *N* = 49 subjects (‘rs-only’) had ages uniformly distributed over the 18-80y range. The second set of *N* = 36 subjects (‘rs+tasks’) was further split into a first (*N* = 15, 20-25 yrs) and a second (*N* = 21, 59-70 yrs) age groups. These two subsets of subjects were scanned in the framework of initially independent studies (using the same MR-scanner) and not initially intended to study dFC, but could be combined at least for subsets of the analyses, to increase sample size whenever possible.

All subjects had no self-reported neurological, psychiatric, or somatic conditions. Subjects in the ‘rs+tasks’ subset, in addition to resting state analyses, were tested behaviourally with a visuomotor task (see later). Furthermore, for 20 of them we also assessed their general cognitive function with the Montreal Cognitive Assessment (MoCA) (Nasreddine et al., 2005). For all the rs analyses of Figs. 1-5 and S1-4, in which cognitive performance was not relevant, we merged the two subsets of subjects. We distinguished for inter-group comparisons between a *‘Young group’* composed of subjects with lower than median age (overall *N* = 42, 18-42 yrs, median age = 24 yrs), and an *‘Older group’* composed of subjects with larger than median age (overall *N* = 42, 47-80 yrs, median age = 63 yrs).

**Fig. 1.**
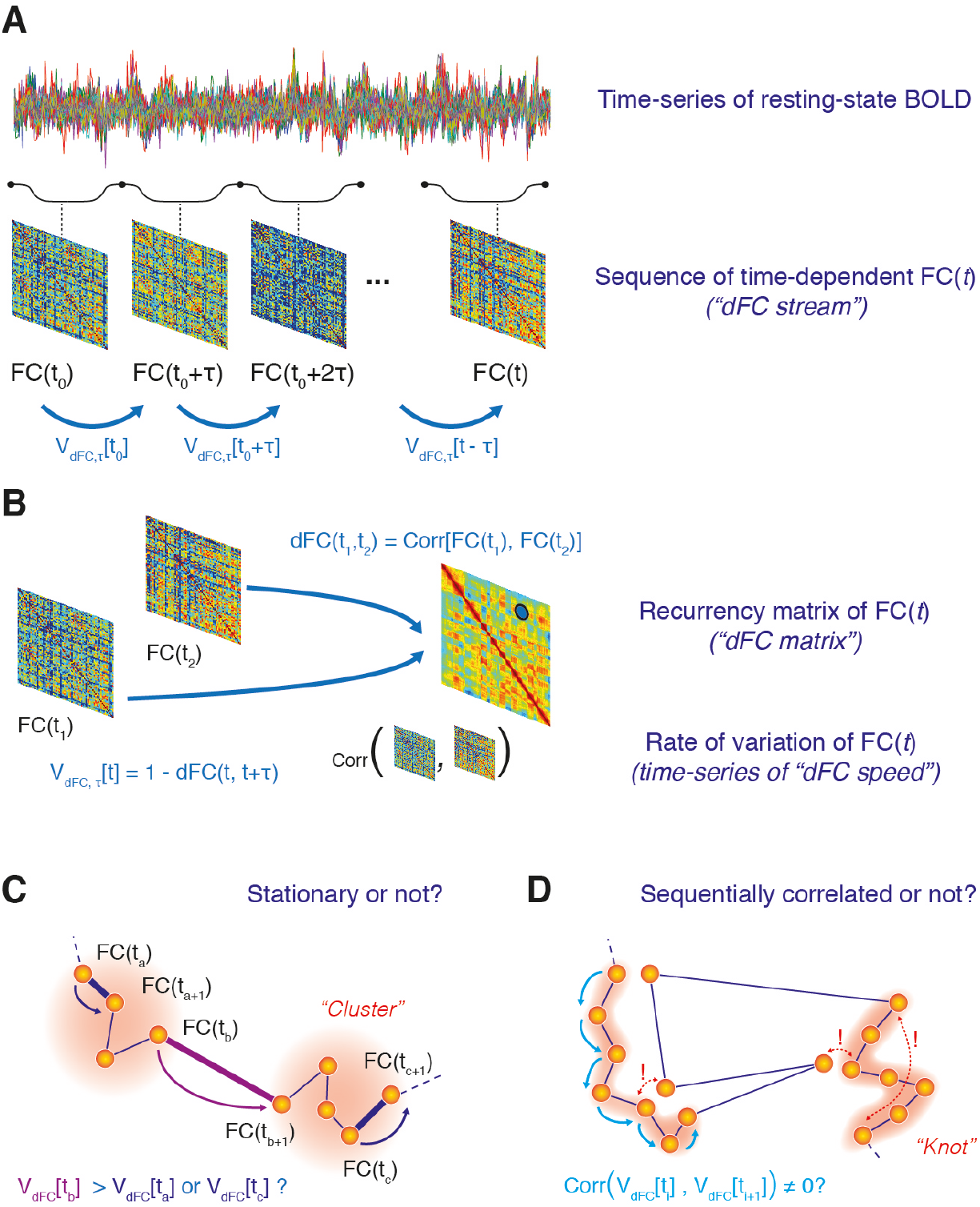
Streams of dynamic Functional Connectivity (dFC). (A) We adopt a sliding-window approach to estimate temporal changes of dynamic Functional Connectivity from fMRI data (resting-state or task). We call the resulting smooth sequence of time-resolved FC*(t)* matrices a *dFC stream*. We then measure at any time *t* the *dFC speeds V_dFC, τ_ (t)* as the variation of FC*(t)* observed between a time *t* and a time *t* + *τ*, where *τ* is the size of the window used to estimate FC*(t)* matrices. (B) The degree of similarity (inter-matrix correlation) between FC*(t)* networks observed at different times can then represented into a recurrence matrix, or *dFC matrix*, whose entry dFC*(t_1_, t_2_)* reports the correlation distance between the functional networks FC*(t_1_)* and FC*(t_2_)* estimated respectively at times *t_1_* or *t_2_*. The block structure of dFC matrices reflects the inhomogeneous speed of variability of FC*(t)* along the flow of the dFC stream. The quantification of dFC speeds allows answering different questions about the statistical properties of dFC streams, summarized by two graphical cartoons: first, (C) whether FC*(t)* matrices tend to cluster into FC states dissimilar between them, but internally similar (stationarity or non-stationarity of dFC); second, (D) whether there are sequential correlations in the dFC stream (e.g. short “jumps” followed by other short “jumps” with high probability), which could be possible even in the case of stationarity. In panel (D), exclamation marks highlight that *“dFC knots”* –epochs of sequentially correlated short jumps– are not clusters since they can have an internal variance larger than the variance between time-resolved FC*(t)* networks separated by *“dFC leaps”*.

**Fig. 2.**
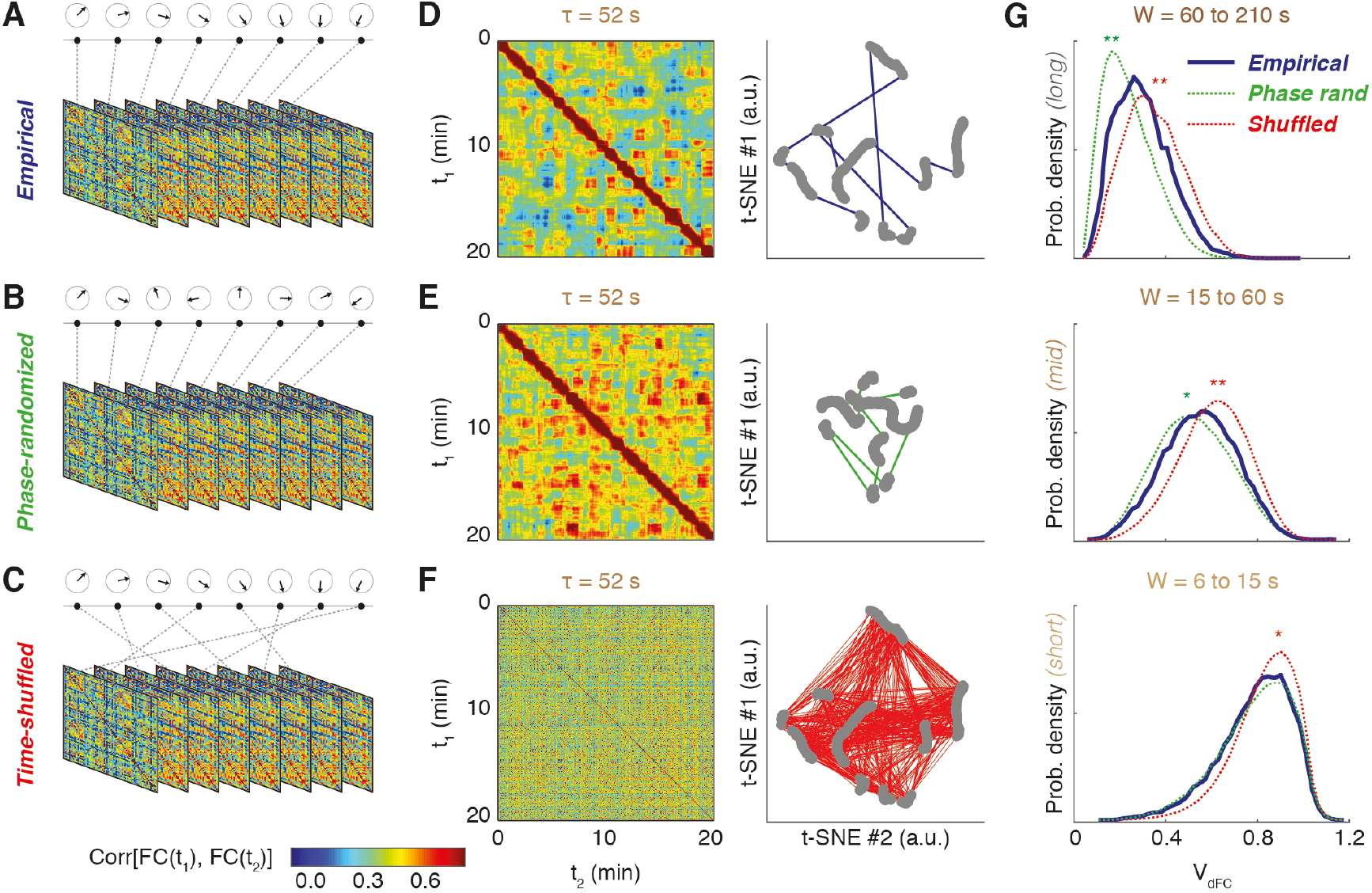
Empirical vs surrogate dFC streams. We compare dFC streams evaluated from actual empirical fMRI data (A) with surrogate dFC streams evaluated: from phase-randomized BOLD time-series surrogates (B), compatible with a null hypothesis of stationarity of the dFC stream; and from time-shuffled surrogates (C), compatible with an alternative null hypothesis of lack of sequential correlations in the dFC stream. We show in panels D-F representative dFC matrices (for a given window size τ = 52 s) for: (D) empirical data; (E) phase-randomized data; and, (F) time-shuffled data. To the right of the dFC matrices we also show a distance-preserving non-linear projection in two dimensions of the associated dFC stream (using the t-SNE algorithm; every dot corresponds to a specific observations of FC*(t)* and the path connecting the dots indicates the temporal order in which the different networks are sequentially visited). Note that the projections of individual FC*(t)* networks in panels (D) and (F) are identical by construction but visited in different orders. These projections make visually evident the stochastic walk nature of dFC streams. (G) Shown here are distributions (smoothed kernel-density estimator) of resting-state dFC speeds for empirical and surrogate ensembles (averaged over subjects), pooled over three distinct window-size ranges: long windows (top, 60 to 210 s); intermediate windows (middle, 15 to 60 s); and short windows (bottom, 6 to 15 s). Distributions for surrogate data significantly differ at the global level from distributions for empirical data in most cases (differences between empirical and time-shuffled distributions in red, between empirical and phase-randomized in green color; stars denote significant differences under two-sided Kolmogorov-Smirnov statistics: *, *p* < 0.05; **, *p* < 0.01).

**Fig. 3.**
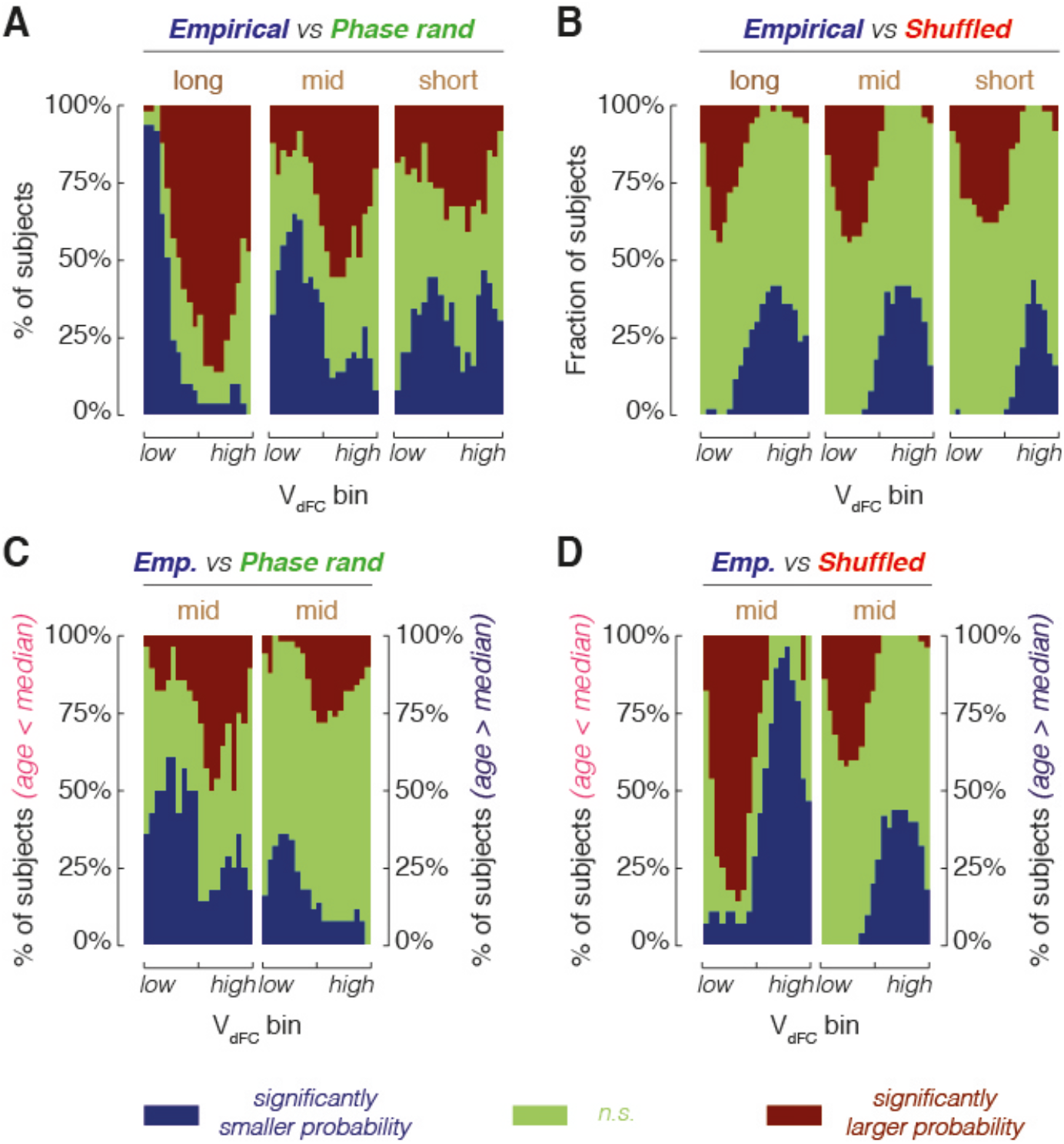
Empirical dFC streams lie “between order and randomness”. We compare empirical and surrogate histograms of resting-state dFC speeds, pooled over three different window-size ranges (long, intermediate, and short) at the single-subject level. We perform comparisons speed bin by speed bin (overall 20 speed bins, ranked from lower to higher speeds), checking whether dFC speeds within a given bin are observed with a probability significantly above chance-level (red), significantly below chance level (blue) or compatibly with chance level (green), according to a selected null hypothesis: stationarity for comparison with phase-randomized surrogates; and lack of sequential correlations for comparison with time-shuffled surrogates. (A) Comparison with phase-randomized surrogates indicates that, in empirical resting-state data, lower (higher) than median dFC speeds are often under-(over-) represented. These effects are particularly evident in the long time-windows range (leftmost plot). (B) Comparison with time-shuffled surrogates indicates that, in empirical resting-state data, lower (higher) than median dFC speeds are often over-(under-) represented, i.e. a reverse pattern relative to phase-randomized surrogates. (C-D) When separating subjects into two age groups (younger or older than the median), the comparison patterns revealed by panels A and B are confirmed, but crisper for young subjects and more blurred for older subjects. Overall, if we dub as “order” the null hypothesis of static average FC (i.e. phase-randomization) and as “randomness” the null hypothesis of temporally uncorrelated dFC fluctuations (i.e. time shuffling), the statistics of resting-state dFC fluctuations appear to lie “between order and randomness” (i.e., they are “complex”).

**Fig. 4.**
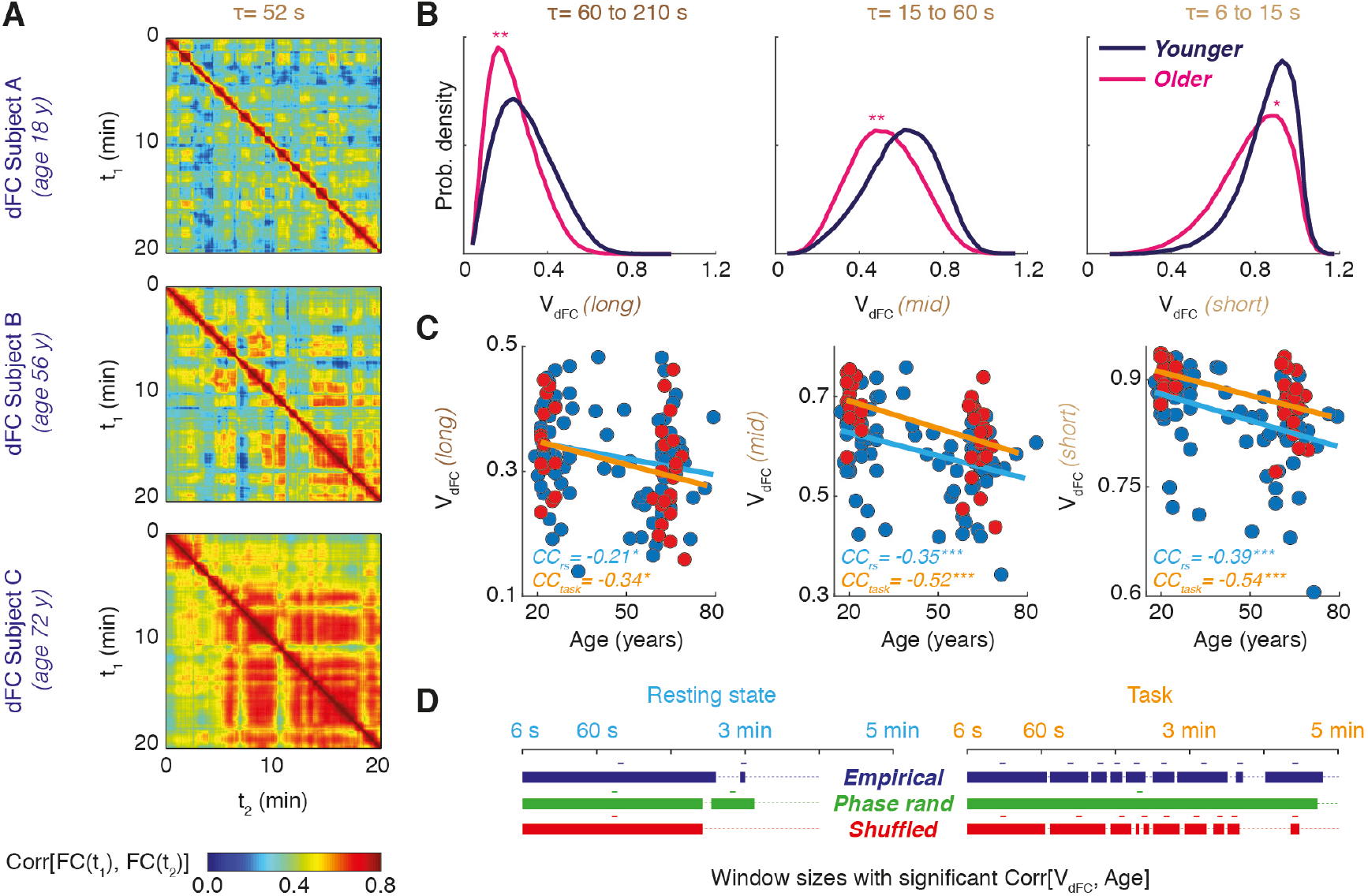
The speed of dFC streams slows down with aging. (A) We show here representative resting-state dFC matrices for three subjects of different ages. With growing age (top to bottom), red block associated to dFC “knots” become longer-lasting. (B) Correspondingly, the distributions of dFC speeds, over all three studied ranges of window sizes, are shifted toward lower values for older relative to younger subjects (one-sided Kolmogorov-Smirnov statistics: *, *p* < 0.05; **, *p* < 0.01). At the single-subject level, median dFC speeds significantly correlate with subject’s age. (C) Scatter plots of single subject age vs median dFC speeds (empirical data), for three pooled window size ranges (long to short, from left to right) and for both resting-state (blue dots) and task (red dots) fMRI scan blocks. Significant correlations with age are found for all three window ranges and for both rest and task dFC speed analyses (bootstrap with replacement confidence intervals for Pearson correlation: *, *p* < 0.05; **, *p* < 0.01; ***, *p* < 0.001). (D) Significant age correlations occur robustly as well for single-window dFC speed estimations over wide continuous ranges of window sizes. We report ranges of window sizes in which statistically significant correlations between median dFC speed and age are detected (a “-” sign indicates negative correlation). To the left, resting-state; to the right, task dFC speeds. Significant correlations (bootstrap, *p* < 0.05) are found not only for empirical data but also for both types of surrogate data in widely overlapping window ranges.

**Fig. 5.**
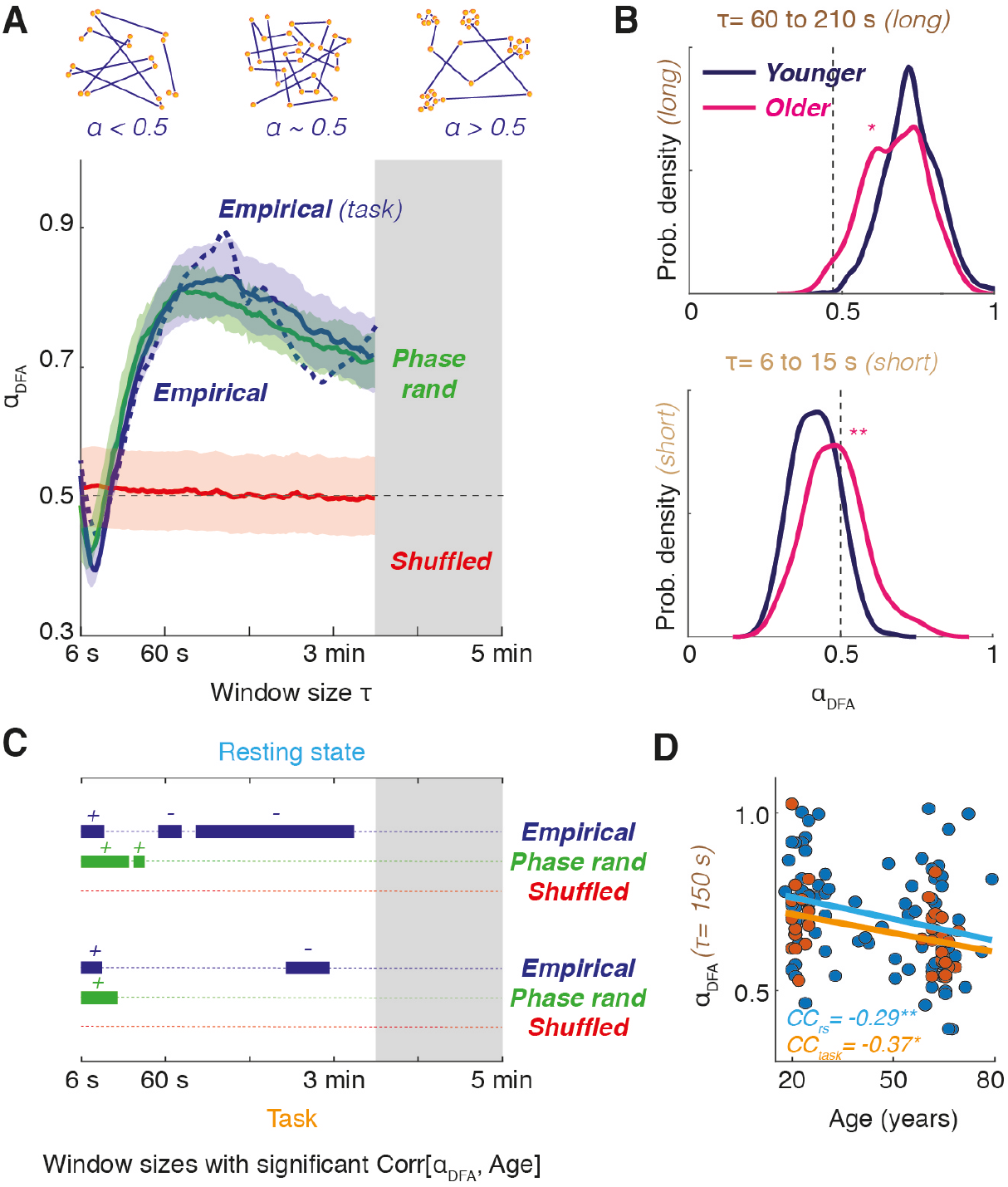
The dFC stream is an anomalous stochastic walk. (A) We performed Detrended Fluctuation Analysis (DFA) of the time-series of instantaneous variations of FC*(t)* along the resting-state and task dFC streams. For every window size, such analysis quantifies whether the stochastic fluctuations of FC*(t)*: are uncorrelated, as in a Gaussian random walk (corresponding to a DFA exponent α_DFA_ ~ 0.5, akin to “white noise”); positively correlated as in a persistent anomalous stochastic walk (corresponding to 0.5 < α_DFA_ < 1, akin to “pink noise”, observed for most intermediate and long window sizes); or negatively correlated as in an anti-persistent anomalous stochastic walk (corresponding to α_DFA_ < 0.5, akin to “blue noise”, observed for short window sizes). Cartoons of these three types of stochastic walk are represented on the top of the plot. Lines report the median α_DFA_ and shaded contours the 95% confidence interval over subjects (median ± 1.96*standard deviation of sample mean). The max τ used for single-window α_DFA_ calculation was shorter than for dFC speed analyses and trimmed to 210 s (excluded τ range shaded in gray). (B) Distributions of resting-state α_DFA_, pooled over all subjects and window sizes within distinct ranges (long windows on top and short windows on bottom) are shifted from persistence or anti-persistence toward the uncorrelated randomness value α_DFA_ ~ 0.5 for older relative to younger subjects (one-sided Kolmogorov-Smirnov statistics: *, *p* < 0.05; **, *p* < 0.01). (C) Significant correlations (bootstrap, *p* < 0.05) between α_DFA_ and age occur over selected ranges of window sizes (a “-” or “+” sign indicate negative or positive correlations). On top, resting state; bottom, task dFC speeds. Robust negative correlations are found only for empirical data. (D) Scatter plots of age vs α_DFA_ (empirical data), for a representative τ in the long window range and for both resting state (blue dots) and task (red dots) fMRI scan blocks.

In addition to general exclusion criteria for participation in an MRI experiment, we excluded subjects with a self-reported musical background, as musical training may affect the performance of rhythmic visuomotor tasks. Left-handed subjects, identified using the Edinburgh Handedness Inventory, were also excluded. Subjects were informed of the procedure of the study and basics of fMRI acquisition, and written consent was obtained before data collection. The studies were performed in accordance with the local medical ethics committee protocol at the Charité Hospital (Berlin, Germany).

### MRI acquisition

Magnetic resonance imaging (MRI) acquisition was performed on a 3T Siemens Tim Trio scanner. Every subject was scanned in a session that included a localizer sequence (3, 8mm slices, repetition time [TR] = 20 ms, echo time [TE] = 5 ms, voxel size = 1.9×1.5×8.0 mm, flip angle [FA] = 40°, field of view [FoV] = 280 mm, 192 mm matrix), a T1-weighted high-resolution image (MPRAGE sequence, 192, 1mm sagittal slices, voxel size 1×1×1mm, TR = 1940 ms, TE = 2.52 ms, FA = 9°, FoV = 256 mm, 256 mm matrix), a T2 weighted image (2:16 minutes, 48, 3mm slices, voxel size 0.9×0.9×3mm, TR = 2640 ms, TE1 = 11 ms, TE2 = 89 ms, FoV 220 mm, 256 mm matrix), followed by diffusion weighted imaging (61, 2mm transversal slices, voxel size =2.3×2.3×2.3 mm, TR = 7500, TE = 86 ms, FoV 220 mm, 96 mm matrix). Subjects were then removed from the scanner to have their EEG cap put on, and then simultaneous fMRI-EEG images were acquired in a single run (BOLD T2*weighted, 32, 3mm transversal slices, voxel size = 3×3×3 mm, TR = 1940 ms, TE = 30ms, FA = 78°, FoV = 192 mm, 64 mm matrix). Five dummy scans were automatically discarded by the Siemens scanner.

During resting-state scans, subjects were to remain awake and reduce head movement. Head cushions served to minimize head movement, and earplugs were provided. Scans for the ‘rs-only’ and the ‘rs+task’ subsets of subjects had different durations. For the ‘rs-only’ subset, 20 min of uninterrupted rs scan were performed. For the ‘rs+task’ subset, 5 min of rs were collected before 20 min of task acquisition (see later), and then further 5 min after the task. We ignored differences between the two rs blocks by concatenating them prior to subsequent analysis.

### fMRI processing

fMRI data were pre-processed following Schirner et al. (2015). Here, FEAT (fMRI Expert Analysis Tool) first-level analysis from the FMRIB (Functional MRI of the brain) software was used. Motion correction was performed using EPI field-map distortion correction, BET brain extraction, and high-pass filtering (100s) to correct for baseline signal drift, MCFLIRT to correct for head movement across the trial. As an additional correction measure, we further regressed out six FSL head motion parameters from the measured BOLD time-series. Functional data was registered to individual high-resolution T1-weighted images using linear FLIRT, followed by nonlinear FNIRT registration to Montreal Neurological Institute MNI152 standard space. Voxel-level BOLD time series were reduced to 68 different brain region-averaged time series, according to a Desikan parcellation (Desikan et al., 2006). See Table S1 for the regions of interest. We neither performed a slice-timing correction, smoothing, normalization of BOLD intensities to a mean, nor global regression.

### Visuomotor coordination task

The *N* = 36 subjects in the ‘rs+tasks’ subset performed a visuomotor coordination task while in the scanner (Chettouf et al., 2020). The task followed a unimanual paradigm, which was adapted from a bimanual paradigm introduced in (Houweling et al., 2008). During the task, subjects were told to lay still inside the scanner with an air-filled rubber ball in their right hand. A screen, animated using a custom-made LabView program, was projected in the scanner. To reduce eye movement, subjects were instructed to fix their gaze at a cross, displayed in the middle of the screen between two rotating disks. The left disk served as a visual cue, rotating at a computer-generated speed, while the subject’s squeezing of the ball controlled the speed of the right disk. The goal was to make the subject-generated rotating disk align in (counter) rotation with the computer-generated rotating disk, which was done by squeezing the rubber ball in a 4:3 frequency to the visual cue. For perfect performance, the two disks would rotate in synchrony. Because the computer-generated disk rotated at a 4:3 frequency to the subject-generated circle, subjects had to squeeze the ball at 1.35 cycles per second to match the 1.8 cycles per second of the computer-generated disk to achieve synchrony

Behavioural measures were collected (one performance score per trial) based on the frequency locking of the two rotating circles. If the two disks rotated perfectly in-synchrony (i.e., subject was able to match the frequency of bulb-squeezing to the computer-generated cue), the performance score would be 1. Not frequency-locked rotations of the two disks would result in a performance score of 0. More specifically, the frequency locking of the computer-generated circle and the subject-generated disk was quantified by the *Spectral overlap* (***SO***) between the power spectra of the two forces, *P_x_* and *P_y_*, as described in detail in (Daffertshofer et al., 2000):

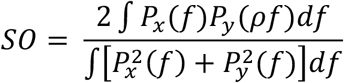

with ρ = 4/3, corresponding to the target frequency ratio between the two rotating disks. Behavioural performance was expected to improve across trials as subjects learned the task; here we focused ‘only’ on the average performance over trials, ignoring all learning effects.

### Cognitive assessment

MoCA assessment was performed by *N* = 21 elderly subjects of the ‘rs+tasks’ subset. The MoCA includes multiple sub-tasks probing different cognitive domains such as: short-term memory and delayed recall; visuospatial abilities; phonemic fluency, verbal abstraction and naming; sustained attention and concentration; working memory; executive control in task switching; spatio-temporal orientation. The test was administered in a German version (downloadable from http://www.mocatest.org). The maximum global score ‘MoCA’ achievable is of 30 points, up to 5 of which are contributed from the partial score ‘MoCA-wm’ from the working memory (‘Erinnerung’) task. Participants were considered in good/healthy mental state, when achieving scores higher than 25. All details can be found in (Nasreddine et al., 2005).

### Control resting state fMRI dataset

To verify that the main results of our analyses did not apply just to our specific dataset, we also repeated some of the analyses on an independent control dataset, selected from rs-fMRI data released as part of the Human Connectome Project (HCP), WU-Minn Consortium. We used the same selected subjects used for statistical benchmarking of functional network analyses in Termenon et al. (2016). This sample includes 99 young healthy adults from 20 to 35 years old (54 females). Each subject underwent two rs-fMRI acquisitions on different days and here, not being interested in test-retesting issue, we used only data from the first day sessions. For this sessions, TR = 720 ms and resting state scan duration was of 14 min and 24 s. More details on data acquisition and pre-processing for this control dataset can be found on Termenon et al. (2016).

### Extraction of time-dependent Functional Connectivity and dFC matrices

In brief, we estimated the sequence of time-dependent Functional Connectivity matrices ***FC(t)*** –or *dFC stream*– by sliding a temporal window of fixed duration ***τ*** (cf. Allen et al., 2012) and by evaluating zero-lag Pearson correlations between resting-state BOLD time series from different brain regions ***i*** and ***j***:

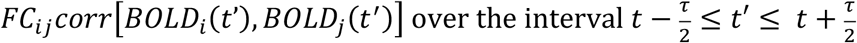

All entries were retained in the matrix, independently of whether the correlation values were significant or not or without fixing any threshold (i.e., we treated ***FC***_***ij***_ entries as descriptive features operationally defined by the above formula).

To evaluate the dFC matrices of Figures 2, 4 and S2 we introduced a notion of similarity between ***FC(t)*** matrices following (Hansen et al., 2015), based on the Pearson correlation between the entries of their upper-triangular parts:

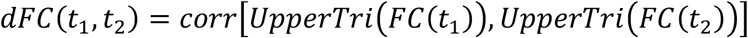

The dFC matrices thus depend on the window-size ***τ*** adopted when evaluating the dFC stream. To perform the two-dimensional projections of the sequence of ***FC(t)*** matrices in Figure 2, the vectors ***UpperTri(FC(t))*** served as input features into a t-Stochastic Neighbourhood Embedding algorithm (exact method) as described by Hinton & Van der Maaten (2008), with default perplexity = 30 and exaggeration = 4 parameters in the employed MATLAB® (MathWorks R2017b) implementation. To compare projections of actual empirical dFC and dFC evaluated for surrogate data (see below) we determined a common projection based on a unified training set combining empirical and surrogate ***FC(t)*** matrices.

### Surrogate dFC models

We compared dFC streams between actual empirical data and two different types of surrogate, to probe deviations from two alternative null hypotheses.

We constructed *phase-randomized surrogates* following Hindriks et al. (2016). In brief, the multi-variate time-series of BOLD signals were first Fourier transformed, extracting time-dependent amplitude and phases at different frequencies for every region. Subsequently, the Fourier phases were drawn from a uniform distribution ***𝒰***(0,2***π***) before applying an inverse Fourier transform yielding surrogate time-series (Kantz & Schreiber, 2004; chap. 7.1.2) with unaltered power spectra. Note, however, that additional care needs to be taken to preserve the overall covariance matrix, as required for a null hypothesis of stationarity. Hence, we used the same random sequence of phases for all the time series, i.e. all the regions, per epoch. We adopted the MATLAB code kindly provided by Rikkert Hindriks and incorporated it in our downloadable dFC toolbox. This approach guarantees that the covariance matrix of the original empirical data is maintained but destroys systematic deviations from stationarity. The phase-randomized surrogates are thus compatible with a null hypothesis of stationary inter-regional FC. Once phase-randomized BOLD time-series have been generated, the dFC stream can be constructed based on them as for the original data.

We then generated *time-shuffled surrogates*. After computing a dFC stream on the actual empirical data, we generated a time-shuffled version by randomly permuting the order of the ***FC(t)*** timeframes but maintaining them individually unchanged. By this, the mean and variance of each of the FC connections independently are preserved but any sequential correlations are disrupted. Time-shuffled surrogates are thus compatible with a null hypothesis of absence of sequential correlations in the dFC stream.

When presenting results for surrogate ensemble we usually generated 1,000 different random realizations of each of the surrogate types for each subject and present average results or over these different realizations, unless differently specified.

### Analysis of dFC speeds

Using correlation as a measure of similarity between matrices implied to use the correlation distance between two observations of functional connectivity as a measure of the amount of change between two ***FC(t)*** observations. By measuring the distance between two FC observations separated by a fixed amount of time set to be equal to the window-size W we thus defined the instantaneous global dFC speed as:

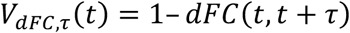

We refer to this dFC speed as “global” because it is evaluated by comparing whole-brain FC matrices. Note that by this definition hte dFC speed depends on the chosen window size ***τ***. We improved the dFC speed histogram estimates by increasing the number of sampled speed observations and avoided potential aliasing artefacts due to the use of a single window (Leonardi & Van de Ville, 2015) by pooling window sizes, given that speed distributions for close enough ***τ*** were similar. We could realise that for a vast majority of subjects and for most dFC speed bins, as binned counts in the histograms extracted at contiguous window sizes were statistically indistinguishable. For 76 out of the 85 included subjects, the binned dFC speed counts in the histograms extracted at window sizes ***τ***_***k***_ and ***τ***_***k***_ + 1 were statistically indistinguishable (overlapping confidence intervals, Agresti-Coull estimation) for at least 12 out of 20 bins, where ***τ***_***k***_ denote the window sizes included (this holds for any of the considered ***τ***_***k***_-s, ranging from 3 to 150 TRs for rs scans in the ‘rs-only’ subset or HCP control dataset and task scans in the ‘rs+task’ subset and from 3 to 105 TR in the –shorter– resting-state scans for the ‘rs+task’ subset). Given this substantial degree of redundancy between speed distributions for contiguous window sizes, we chose to pool speed samples to form just three histograms, over three (arbitrary) window ranges: a “short” window-size range, ~6 < ***τ*** < ~15 s (3 to 8 TRs); a “mid” window-size range, ~15 s < ***τ*** < ~60 s (9 to 32 TRs); and a “long” window-size range, ~60 s < ***τ*** < ~210 s (33 to 105 TRs). Window pooling leads to smoother dFC speed histograms. The three different ranges were chosen to construct in each range histograms with close numbers of speed observations after pooling (note that for longer windows less speeds are computed because there are less non-overlapping windows pairs, so that more window sizes must be pooled to reach a number of observations close to the short windows range). A similar window-pooling strategy is used as well in our related study by Lombardo et al. (2020), where we also introduce metrics of dFC speed restricted to specific networks of interest.

Correlations and scatter plots between age or cognitive scores and dFC speeds were constructed based on the median of dFC speed distributions, either computed at single window sizes or pooled, depending on the different analyses. The same procedures were followed for all dFC speed analyses for the actual empirical and for the two types of surrogate data. For comparing dFC speed histograms between empirical and surrogate data (as in Figures 3 and S3, S4 or S5), we used a same binning in 20 bins for the three types of data for every subject, but adapted the binning to each specific subject’s range, always keeping the same number of bins. Next, we compared subject-by-subject the normalized counts between empirical and surrogate histograms, proceeding from the leftmost to the rightmost bin. Finally, we tested in how many subjects the dFC counts – bin-by-bin – for empirical data were under- or over-estimated (lack of overlap between 95% Coull-Agresti confidence intervals for the count) with respect to histograms for a given surrogate type.

### Detrended Fluctuation Analysis

Detrended Fluctuation Analysis (DFA) allows for detecting intrinsic statistical self-similarity embedded in a time series. It is particularly adapted to the study of time series that display long-range persistence, and it is in this sense similar to other techniques, such as Hurst exponent analysis, the latter requiring however the stationarity of the analysed signal. See Witt & Malamud (2013) or Metzler et al. (2014) for a review. DFA infers a self-similarity coefficient by comparing the detrended mean square fluctuations of the integrated signal over a range of observation scales in a log-log plot. If the log-log plot has an extended linear section, (i.e. if the scaling relation is a genuine power-law over a reasonably broad and continuous range of scales, see later for the meaning of ‘genuine’), it means that fluctuations ‘look the same’ across different temporal scales, i.e. we have statistically the same fluctuations if we scale the intensity of the signal respecting the DFA exponent.

To perform DFA we first evaluated dFC streams (using a given window size τ) and then evaluated its *instantaneous dFC increments*:

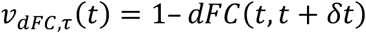

where ***δt*** corresponds here to the minimum possible sliding time-shift in our discretely sampled time-series, i.e. 1 TR (one data point of shift). Note that these instantaneous increments ***ν***_***dFC,τ***_***(t)*** continue to depend on the window-size ***τ*** because the dFC streams are computed adopting a specific window-size. Therefore, we perform a different fractal scaling analysis over each of the dFC streams evaluated for different window sizes ***τ***.

To perform DFA, we first converted the time-series of instantaneous dFC increments ***ν***_***dFC,τ***_***(t)*** into an unbounded process:

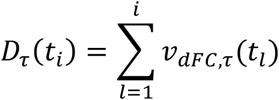

Let ***K*** denote the number of samples in the time series, that are split into ***M*** non-overlapping segments ***q = 1 … M*** of length ***k*** each, with ***M = ⌊K/k⌋***. For each segment *q* the fluctuation strength was computed as the squared difference between ***D_τ_(t)*** and its trend 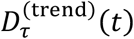 (in the linear case this is the regression line of ***D_τ_(t)*** over the interval ***t = 1 … k***):

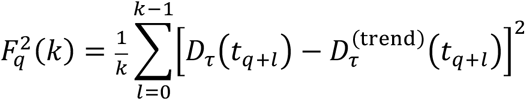

In the case of scale-free correlation this fluctuation strength scales with segment size ***k***. That is, (on average) one finds a linear power law of the form:

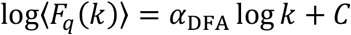

In figure S6 we denote ***F_DFA_(k)*** = **〈*F_q_(k)*〉**. The scaling parameter *α*_DFA_ is the primary outcome measure of DFA. In the case of the scale-free processes with the aforementioned power law, ***α_DFA_*** resembles the Hurst exponent (Metzler et al., 2014), leading to the interpretation:

- 0 < ***α_DFA_ < 0.5: ν_dFC,τ_(t)*** displays anti-persistent fluctuations
- ***α_DFA_ = 0.5: ν_dFC,τ_(t)*** displays uncorrelated Gaussian fluctuations (“white noise” or, equivalently, *D_τ_* resembles Brownian motion)
- ***0.5 < α_DFA_ < 1: ν_dFC,τ_(t)*** displays persistent fluctuations (approaching “pink noise” when ***α_DFA_*** is close to 1)
- ***1 ≤ α_DFA_: ν_dFC,τ_(t)*** is non-stationary (strictly speaking, DFA is undefined in this case)

Prior to construing outcome values, however, it is mandatory to verify that a linear power law scaling actually exists. If it was not the case indeed the output value ***α***_***DFA***_ could not be interpreted as a scaling exponent. Following (Ton & Daffertshofer, 2016; see https://github.com/marlow17/FluctuationAnalysis for a dedicated toolbox), we tested the hypothesis of power-law scaling using a (Bayesian) model comparison approach. This allowed identifying the subjects for which the DFA log-log plot was better fitted by a straight line than by any other tested alternative model. Only these subjects with a proper linear section in the DFA log-log plot were retained for the following steps of DFA exponent extraction and analysis of correlations with age.

In order to test the hypothesis of power law against alternative models, we evaluated the density of fluctuations over the consecutive segments, i.e. the density of ***F_q_(k)*** – beyond its mean value **〈*F_q_(k)*〉** *–* using a kernel source density estimator. Based on this probability density, one can estimate the log-likelihood for a certain model to generate fluctuations of a given strength (on a log-scale) as a function of **log *k***. To perform model selection, the toolbox then computes the corrected Akaike Information criterion, **AIC_*c*_**, for each one of the tested models:

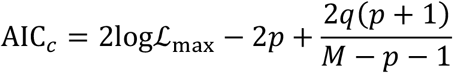

where ***p*** is the number of free parameters to fit in the model and the number of points used to estimate the density of ***F_q_(k)***. Note that this model selection criterion automatically embeds a penalization for models with larger number of parameters, thus protecting against over-fitting. The model yielding the lowest **AIC_*c*_** was selected as the relatively best one, and if this was the linear one, the corresponding 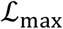-fitting parameter was considered as ***α***_***DFA***_.

We used a range of 10 < ***k*** < 80, to discard data chunk sizes that were too short or long data chunk sizes yielding an overall number ***M*** of chunks that was too small. A genuine power-law scaling in the DFA of subjects could be found for all subjects in the ‘rs-only’ and ‘rs+task’ subsets in at least 80 out of the 105 different window-sizes (3 to 105 TRs) used to estimate the dFC streams. Given this general evidence for widespread power-law scaling of the ***ν***_***dFC,τ***_***(t)*** increments in all subjects, during both resting-state and task scans (apart from sporadic exceptions), as well as in the HCP control data, we computed ***α***_***DFA***_ exponents in all cases.

## Results

### Dynamic Functional Connectivity as a stochastic walk

A widespread manner to extract FC from rs- or task-fMRI is to parcellate the brain into ***N*** macroscopic regions – we follow here a parcellation described first by Desikan et al. (2006) (see Table S1) – and to compute pairwise linear correlations ***FC_ij_ = *Corr*[a_i_(t), a_j_(t)]***, between the region-averaged time series of neural activity ***a_i_(t)*** and ***a_j_(t)*** of regions ***i*** and ***j***, based on the entire fMRI imaging session (tens of minutes). The result of this procedure is a square ***N***-times-***N*** matrix of *static FC*. To estimate how FC fluctuates in time around the session averaged static average, we adopted a common sliding window approach – followed e.g., by Allen et al. (2012) –, repeating the FC construction separately for each time-window (of a fixed duration *τ*) and generated a time-ordered sequence of ***FC(t)*** matrices, or *dFC stream* (Fig. 1A).

Next, we studied the similarity between time-resolved networks observed at different times. To quantify the amount of FC network variation, we introduced a metric of similarity between FC matrices and evaluated the so-called *dynamic Functional Connectivity (dFC) matrices* (cf. Hansen et al., 2015). The dFC matrix entries ***dFC_ab_*** provide the normalized correlation ***Corr[FC(t_a_), FC(t_b_)]*** between any two ***FC(t)*** networks observed at times ***t_a_*** and ***t_b_***, as depicted in Fig. 1B). A rs-dFC stream for a representative resting-state recording is shown in Fig. 2A, with its associated dFC matrix in Fig. 2B (to the left). From the inspection of this matrix, we can recognize that the rate of variation of ***FC(t)*** matrices was not constant along time, but rather heterogeneous. The associated dFC matrix of Fig. 2B (left) displayed characteristic patterns composed out of square-shaped red-hued blocks, corresponding to epochs of transiently increased similarity between consecutive ***FC(t)*** network frames. Such epochs of relative FC stability increase –or ‘*dFC knots*’– were intertwined with transients of relative instability, shown in Fig. 2B (left) by light green or blue stripes in the dFC matrix, denoting strong dissimilarity from previously visited FC*(t)* networks. During such transients –or ‘*dFC leaps*’– ***FC(t)*** quickly morphed before stabilizing again into the next dFC knot. Knots and leaps could be observed when computing dFC streams and matrices over the whole broad range of window-sizes we tried (between 6 s to 5 mins). Additional examples for representative subjects and window sizes are shown in Fig. S2A.

To provide a more quantitative description of the heterogeneous ***FC(t)*** change, we determined the rate of change of FC networks along the dFC stream. As said, we considered the dynamics of FC as a stochastic exploration of the space of possible FC configurations and assumed pairs of ***FC(t)*** matrices to be separated by an observation window equal to the window-size itself *τ* used for ***FC(t)*** estimation. We defined the *global dFC speed* at time *t* as the quantity ***V_dFC,τ_(t) = 1 − Corr[FC(t), FC(t + τ)]***. As illustrated on the bottom of Fig. 1A, this global dFC speed can be interpreted as the distance travelled in FC space between two ‘stroboscopic’ observations at times ***t*** and ***t + τ*** (i.e. over a fixed time interval corresponding to the closest possible interval separating two windows without overlap). Hence, the time-resolved ***FC(t)*** matrix may be seen as performing a stochastic walk in the space of possible FC network configurations. The global dFC speed ***V_dFC,τ_(t)*** thus informs us about how fast and far away is the time-resolved FC network moving along the stochastic path given by the observed dFC stream, at a time ***t***.

We sampled the statistical distributions of ***V_dFC,τ_(t)*** for different subjects and values of *τ* (see *Materials and Methods*). We also computed window-pooled dFC speed, simultaneously mixing estimations from different window sizes within three different ranges –long (one to three minutes), intermediate (tens of seconds) and short (6-15 s) window sizes–, also to avoid detection of artefactual fluctuations due to the use of a unique fixed window (Leonardi & Van de Ville, 2015). A distribution of resting-state pooled dFC speeds over all subjects is shown by the blue curve in Fig. 2C, for the three long, intermediate and short window-size ranges (from top to bottom). Analogously, examples of pooled dFC distributions sampled for single subjects are given in Fig. S2B. For all considered *τ*-s (both at the group and at the single subject level), these global dFC speed distributions displayed a clear peak near a median value ***V_dFC_***, i.e. the *typical dFC speed*. The dFC speed distributions were generally skewed and/or kurtotic, deviating from Gaussianity (over three quarters of computed distributions, for different subjects and window sizes, Lilliefors test, *p* < 0.05) and had generally fat tails, in particular evident for long or short window sizes. Before we discuss the implications of these deviations from Gaussianity, we here want to highlight these fat tails indicate that FC reconfiguration events of an anomalously small (low dFC speed) or large (high dFC speed) size are observed with anomalously large probability.

Overall, the FC reconfiguration speed along the dFC stream appeared not to be constant. To show this, we performed a non-linear distance-preserving projection of the sequences of ***FC(t)*** matrices observed along the dFC stream into a space of lower dimension. We used two-dimensional projections of dFC streams via t-Stochastic Neighborhood Embedding (t-SNE, Hinton & van der Maaten, 2008). In these plots, each dot corresponds to the projection in two dimensions of a different time-resolved FC network and temporally consecutive dots are linked by a line (see *Materials and Methods*). The dFC stochastic walk can thus be explicitly visualized, as in the example of Fig. 2B (right), showing the projection of the dFC stream associated to the dFC matrix plotted on the left. In this t-SNE projection, ***FC(t)*** within dFC knots form smooth and continuous segments, interrupted by a few cusp points, associated instead to dFC leap events. Here we would like to note that that dFC speed fluctuations did not reflect mere head motion artefacts because the size of instantaneous dFC variations (estimated using two alternative forms of motion correction, cf. *Materials and Methods*) did not correlate with the size of instantaneous head displacements; see Fig. S1A. After motion correction, large dFC speeds could still be detected even in absence of head movement. Conversely, large head movements could occur without big changes of FC. Therefore, dFC fluctuations are most likely not artefactual. We would also like to note, that the very fact that dFC speed varied does not readily imply that dFC stream to be trivial: To assess “non-triviality”, one must be able to prove deviations of the statistical properties of empirical dFC streams from trivial null hypotheses. Below we discuss possible deviations from two alternative null hypotheses of “order” and “randomness”. To anticipate, empirical dFC streams have dFC speed distributions lying between these two null hypotheses (Crutchfield, 2011; see *Discussion*).

### Dynamic Functional Connectivity deviates from “order”

***FC(t)*** variability might not be indicative of genuine dynamics but represent fluctuations around an underlying “order” described by an unchanging, static FC. This “order” scenario corresponds to a null hypothesis of *FC stationarity*. In contrast to this is the possibility that a multiplicity of separate FC states exists. As represented in the graphical cartoon of Fig. 1C, deviations from stationarity would imply that well-defined clusters of ***FC(t)*** matrices exist along the stochastic walk described by the dFC stream. The step lengths travelled in FC space –measured by dFC speed– would thus be significantly shorter between ***FC(t)*** matrices belonging to a same cluster than the distance travelled between ***FC(t)*** matrices belonging to different clusters. A possibility is thus that at least some of the red-hued “knots” observed in dFC matrices (cf. Fig. 2B) correspond to well-separated clusters, associated with distinct FC states. To test for this possibility, we generated *phase-randomized* surrogate dFC streams, following Hindriks et al. (2016). In this type of surrogates (Fig. 2D), a phase-shift –randomized at every timepoint but identical for all regions– is applied to fMRI time-series, such to preserve by construction the static time-averaged FC of the measured session but to destroy any coherent fluctuations around it that may result in deviations from stationarity (see Materials and Methods and Hindriks et al. (2016)). When computing dFC streams, dFC matrices and dFC speed distributions for phase-randomized surrogates, we found that ***FC(t)*** fluctuations were not suppressed. Phase-randomized dFC matrices still appeared to have blocks and stripes reminiscent of “knots” and “leaps” in empirical dFC streams (Fig. 2E, left). Analogously, when performing a low-dimensional projection via the t-SNE algorithm, phase-randomized dFC streams still gave rise to stochastic exploration paths alternating continuous sections with discrete jumps (Fig. 2E, right). The observed larger-than-average fluctuations of ***FC(t)*** were well compatible with the null hypothesis of stationarity. However, when computing the distributions of dFC speed for phase-randomized surrogates, we found that they generally differed from empirical distributions, both at the group level and at the level of single subjects. The green dashed curves in Fig. 2C show distributions of resting-state pooled dFC speeds over phase-randomized surrogates for all subjects, for the three long, intermediate and short window-size ranges (from top to bottom). Once again, these distributions were skewed and/or kurtotic, tending to deviate from Gaussianity (Lilliefors test, *p* < 0.01 for long and short windows, not significant for intermediate windows). Most importantly, these pooled dFC speed distributions for phase-randomized surrogates were statistically different from equivalent empirical distributions for two out of three window-size ranges (two-sided Kolmogorov Smirnov test, Bonferroni corrected, *p* < 0.01 for long windows; *p* < 0.05 for intermediate windows; not significant for short windows). The distribution mode was smaller than for the empirical dFC streams.

We performed comparisons at the level of single-subject dFC speed histograms. As shown in Fig. 3A, the probability of observing speeds in the slow range was significantly smaller in empirical distributions than in phase-randomized distributions for a majority of subjects. At the same time, the probability of observing speeds in the fast range was significantly larger (95% binomial confidence interval speed bin-by-speed bin comparison). We obtained similar results when analyzing single subject distributions of dFC speeds during task blocks, rather than resting state (Fig. S3A). Hence, both at the group and single-subject levels, phase-randomized dFC speed distributions appeared shifted *toward slower speeds*, with respect to empirical dFC streams. Note that this deviation of empirical dFC speed distributions from the stationary “order” scenario still does not imply that red-hued “knots” in the dFC matrix are actual clusters and that “leaps” are FC state transitions. Indeed these “knots” and “leaps” are visible even in dFC matrices for stationary phase-randomized surrogates (Fig. 2D), as previously mentioned. Furthermore, by construction, phase-randomized surrogates must yield the same covariance matrix of empirical data and, therefore, the level of ***FC(t)*** variability for empirical and phase-randomized data should be the same. How can these observations be reconciled with a statistically significant difference in dFC speed distributions? We will comment on this apparent contradiction in the *Discussion*. However, for now, we would like to remark that deviations from the stationary order expressed by the phase-randomized surrogates do not mean automatically that dFC streams are non-stationary. Still, it means that they are “*differently stationary”*.

We also note that the results obtained for our original dataset hold even for an independent dataset mediated from the Human Connectome Project (HCP, see *Materials and Methods*). Indeed, as visible in Fig. S4A, distributions of dFC speeds for phase-randomized surrogates are shifted toward slower speeds than for empirical control data, both at the group level (Fig. S4A) and at the single-subject level (Fig. S4B) and at all window-size ranges.

### Dynamic Functional Connectivity deviates from “randomness”

The notion of dFC speed does not capture generic variance across ***FC(t)*** matrices observed at any point in time but specifically focus on *sequential variability*, i.e. on variability occurring along a time-ordered dFC stream between a matrix ***FC(t)*** and a second one at a fixed time-distance ***FC(t + τ)***. The null hypothesis of stationarity imposes a static covariance matrix but does not make differences about the “when” relatively larger and smaller fluctuations occur. “Knots” and “leaps” are localized in specific time-epochs. Waxing and waning knots have a beginning, a duration and an ending, depicted as blocks in the dFC matrix. To test for sequential aspects in the empirical dFC streams, we designed a second trivial scenario of maximal sequential “randomness”. In this alternative null hypothesis –associated with *shuffled* surrogate dFC streams–, ***FC(t)*** means and variances are preserved, while sequential correlations along the stream are destroyed by randomly shuffling the order of timeframes in the empirical dFC stream (Fig. 2F). Contrary this scenario is the possibility that “flight lengths” in the space of FC matrices are sequentially correlated. For instance, as sketched in the cartoon of Fig. 1D, short steps may be followed by short steps with a larger than chance probability –a property of the stochastic walk known as *persistence* (Witt & Malamud, 2013; Metzler et al., 2014; see later). If “knot” epochs are sufficiently long-lasting, ***FC(t)*** may evolve considerably –and do so smoothly and very gradually– through the composition of a multiplicity of small steps. In this way, transient slowing downs would not be automatically associated to the emergence of FC clusters (as in Fig. 1C). Distances between ***FC(t)*** matrices visited during a slowing-down epoch could indeed be even larger than distances between matrices visited during different slowed-down transients (compare e.g., the distances marked by an exclamation mark in Fig. 1D). However, slowing down transients and transient accelerations would still appear in the dFC matrix as red-hued “knot” blocks and blueish “leap” stripes. Incidentally, this discussion confirms once again that the appearance of “knots” and “leaps” in the dFC matrix is not proof per se of the existence of FC clusters and states. But are the “knots” indicative of significant transient slowing-downs with respect to chance level set by the “randomness” null hypothesis? The full proof will require a direct, explicit study of the persistence of the dFC stochastic walk (see below). Here, we first study differences at the level of the distribution of dFC speeds, as we already did with the “order” null hypothesis.

In Fig. 2E we show the dFC matrix and the two-dimensional projection associated to a typical shuffled dFC stream. The dFC matrix (Fig. 2E, left) appears powdery and scattered without visible knot and leap patterns. This fact is not surprising, because this matrix contains precisely the same entry values of the original empirical dFC matrix in Fig. 2B but in a randomly permuted order. Analogously, the individual ***FC(t)*** matrices of the shuffled dFC stream are identical to the matrices composing the empirical dFC stream but appear in a permuted order. As a result, in the t-SNE projection of Fig. 2E (right), the points associated with each of the individual ***FC(t)*** timeframes are precisely identical to the ones in Fig. 2B (right). The path linking them is erratic and unstructured, much related to Brownian motion.

When computing the distributions of dFC speed for shuffled surrogates we found that they generally differed from empirical and phase-randomized distributions, both at the group level and at the level of single subjects. The red dashed curves in Fig. 2C show distributions of resting-state pooled dFC speeds over shuffled surrogates for all subjects, for the three long, intermediate and short window-size ranges (from top to bottom). These distributions had a residual skewness but differed significantly from a Gaussian only for short window sizes (Lilliefors test, *p* < 0.05 for short windows, not significant for other windows). Once again, they significantly differed from matching empirical dFC speed distributions in all the window ranges (two-sided Kolmogorov Smirnov, Bonferroni corrected, *p* < 0.01 for long and intermediate windows; *p* < 0.05 for short windows). The distribution modes were larger than for empirical dFC streams. When performing speed bin-by-speed-bin comparisons at the level of single-subject dFC speed histograms, the pattern was reversed with respect to the comparison with phase-randomized surrogates. Notably, as shown in Fig. 3B, the probability of observing speeds in the slow range was significantly larger in empirical distributions than in shuffled distributions for a majority of subjects. At the same time, the probability of observing speeds in the fast range was significantly smaller (95% binomial confidence interval speed bin-by-speed bin comparison). We obtained similar results when analysing single subject distributions of dFC speeds during task blocks, rather than resting state (Fig. S3B).

Both at the group and single-subject levels, shuffled dFC speed distributions appeared shifted *toward faster speeds* relative to empirical dFC streams. Similar results hold as well for the control HCP resting state fMRI dataset, as shown by Fig. S4A (for group-level differences) and S4C (for single subject-level differences). Summarizing, the results of comparisons of the empirical dFC speed distributions with the two alternative types of surrogates, we conclude that, in both rest and task conditions, the empirical dFC speed distribution lies between the trivial “order” and “randomness” scenarios. It is thus reflecting a non-trivial disorder, a.k.a. complexity (see *Discussion*).

### Slowing of dFC through the human adult lifespan

Our fMRI dataset included subjects over a wide age range of 18 to 80 years. The enabled us to study how dFC stream properties are affected by “healthy aging” (none of the subjects were diagnosed with a pathological decline of cognitive abilities). Fig. 4A shows dFC matrices for representative subjects of different ages (see Fig. S2A for different *τ*-s). It is visually evident that the typical duration of dFC knots varied with subject age, seeming to become longer for older subjects.

To quantify this visual impression, we computed in Fig. 4B resting-state pooled dFC distributions separately for the group of subjects younger (blue curves) and older than median age (magenta curves), for the three long, intermediate and short window ranges (from left to right). For all the three window ranges, the distribution of dFC speeds for older subjects was significantly shifted toward slower values (one-sided Kolmogorov-Smirnov, *p* < 0.01 for long and intermediate windows, *p* < 0.05 for short windows), reflecting the longer duration of the slowing-down epochs associated to knots in the dFC matrix. Similar results held for task dFC speed as well.

At the level of single-subject pooled dFC speed distributions, we tracked for every subject the position of the distribution medians, giving the typical dFC speed ***ν_dFC_*** and computed its correlation with subject age over the different window ranges. As shown by the scatter plot in Fig. 4C, ***V_dFC_*** significantly decreased with age (bootstrap with replacement c.i., see caption for *p* values) for all three pooled window ranges, for resting-state and –with even stronger correlations– for task dFC speeds. The correlations of dFC speed with age where robust even when considering single window estimations, without pooling. As visible on the top of Fig. 4D (blue ranges), single window ***V_dFC_***-s correlated negatively with subjects age over a broad range of sizes ranging from very-short window sizes (6-15 s) up to window sizes of several minutes.

We also found that these negative correlation between ***V_dFC_*** and age held as well for both phase-randomized and shuffled surrogates, and over broad window ranges well matching the ones found for empirical data (see green and red ranges in Fig. 4D). This suggests that dFC slowing down may be linked to a general reduction of FC*(t)* variance, which is retained as well when constructing the surrogate dFC streams (see *Discussion*). More specifically, we found that the differences between surrogate and empirical dFC speed distribution tended to smear out with increasing age. In Fig. 3C-D (for the intermediate window range) and in Fig. S5 (for short and long window ranges), we repeated the speed bin -by- speed bin comparison of empirical with surrogate single subject dFC distributions, separating now, however the subjects into two age groups, younger or older than the median age. The patterns of comparison with surrogate distributions have the same overall directions for younger or older subjects: slower speeds tend to be under-represented (over-represented) and faster speeds over-represented (under-represented) when comparing empirical with phase-randomized (shuffled) dFC distributions. However, for the older subjects, the fraction of subjects for which the bin-by-bin comparisons with surrogates were not substantially increased (cf. the broader green central band in Figs. 3C-D or S5). In a sense, therefore, the dFC speed distributions get more “trivial” with aging (i.e., less complex).

### Dynamic Functional Connectivity is an anomalous stochastic walk

As previously said, stochastic processes can be “memory-less” –the next step is uncorrelated from the preceding– or display long-range correlations (Witt & Malamud, 2013; Metzler et al., 2014), positive (*persistence*) or negative (*anti-persistence*). The classic “Drunkard’s walk”, associated with Brownian motion (or white noise), corresponds to a Gaussian, memoryless process. However, other stationary stochastic processes can display long-range sequential correlations resulting in different statistics of fluctuation, toward “pink noise” (for positive correlations and persistence) or “blue noise” (for negative correlations and anti-persistence). Such *anomalous* –i.e., deviating from Gaussianity– are common in a variety of contexts (see *Discussion*) and has been already identified in resting-state and task fluctuations of both electrophysiological and fMRI signals (Linkenkaer-Hansen et al., 2001; Van de Ville et al., 2010; He, 2014).

To characterize deviations from Gaussianity in the FC*(t)* fluctuations along the dFC stream, we used a quantitative approach, Detrended Fluctuation Analysis (DFA, Kantelhardt et al., 2001) to quantify the degree and type of long-range correlations along the dFC stream of ***FC(t)*** networks. The DFA procedure quantifies the strength of auto-correlations in a sequence by detecting a power-law scaling – described by a scaling exponent *α*_DFA_ – in the divergence of a quantity ***F_DFA_(k)***, probing the strength of the fluctuations of the sequence at different scales of observation *k* (see *Material and Methods*). A value of *α*_DFA_ = 0.5 corresponds to a Gaussian white noise process, in which the standard deviation of the fluctuations grows as √*N* after *N* uncorrelated steps. In contrast, larger values 0.5 < *α*_DFA_ ≤ 1 correspond to anomalously persistent and 0 < *α*_DFA_ ≤ 0.5 anomalously anti-persistent fluctuations. Here, specifically, we measured along the dFC stream sequences of instantaneous increments *v_dFC,τ_ (t)* = 1 − dFC*(t*, *t*+*δt)*, where *δt* correspond to the time-step between one FC*(t)* network frame and the following (i.e., 1 TR). This choice yielded a description of the dFC stream as close to continuum in time as possible. We then applied the DFA procedure on sequences of *v_dFC,τ_* supplemented by a (Bayesian) model-selection step (see *Material and Methods)*, which allows for discarding subjects, for which a genuine power-law scaling is not present (Ton & Daffertshofer, 2016). Note that the DFA exponents *α*_DFA_ will continue depending on the window *τ* chosen for FC*(t)* estimation. Indeed, instantaneous increments *v_dFC,τ_ (t)* –analogously to “stroboscopic” dFC speeds *V_dFC,τ_ (t)*– are necessarily evaluated on a given input dFC stream. Therefore, different dFC streams are obtained for different choices of *τ*. It will thus be necessary to study how *α*_DFA_ depends on *τ*, since different fluctuation statistics could be found for the different ways of measuring FC*(t)* (see *Discussion* for possible alternatives and their drawbacks).

We show in Fig. 5 the results of DFA analyses on our data. First of all, we found robust power-law scaling relations in ***ν_dFC,τ_*** fluctuations over most subjects and window sizes. Fig. S6 shows examples of robust power-law scaling under DFA for representative subjects and window sizes ***τ***. We also found that for window sizes ***τ*** ≥ ~20s (i.e., roughly over the intermediate and large windows ranges) the exponent *α*_DFA_ was systematically and significantly (lack of overlap between 95% mean c.i.) *larger* than 0.5, for both empirical resting-state and task data (sample mean of *α*_DFA_ is given by the blue curves in Fig. 5A), indicating persistence of dFC fluctuations. On the contrary, for window sizes ***τ*** ≤ ~15s (i.e., over the short windows range), the exponent ***α***_***DFA***_ was significantly *smaller* than 0.5, denoting anti-persistence. For nearly all window sizes, dFC stream was thus significantly deviating from a Gaussian random walk displaying important long-range correlations. Remarkably, we found the same pattern of deviation from Gaussianity as a function of the chosen window size for the control HCP resting state dataset (see Fig. S4D). In this case, for window sizes ***τ*** ≥ ~100s, the average *α*_DFA_ exponent indicated a persistence even larger than for our aging dataset.

When performing DFA analysis on surrogate dFC streams, we found by construction that the “randomness” shuffled surrogates corresponded for all probed windows to a Gaussian uncorrelated walk with ***α***_***DFA***_ ~0.5 (sample mean given by the red curve in Fig. 5A). Within computational limits, the “order” phase-randomized surrogates (sample mean given by the green curve in Fig. 5A) had an ***α***_***DFA***_ spectrum statistically indistinguishable from the one of empirical data, in agreement with the literature (Dingwell & Cusumano, 2010), and providing therefore a robust benchmark to probe for eventual correlations between ***α***_***DFA***_ and behaviour or cognition.

Note that the *α*_DFA_ exponents measured for dFC streams strongly differ for analogous exponents estimated from head motion time-series (cf. Fig. S7A), hinting at their origin in genuine fluctuations of neuronal activity. Also note that deviations from Gaussianity in fluctuations of dFC streams may be linked to the underlying spectral properties of fMRI multivariate time-series (cf. Ciuciu et al., 2012, 2014; see *Discussion*).

### Loss of complexity of dFC through the human adult lifespan

We then studied how the scaling properties of dFC stream fluctuations varied with age. We show in Fig. 5B sample resting-state distributions of ***α***_***DFA***_, pooled over the long (top) and short (bottom) window size ranges, separately for the two groups of subjects with smaller (blue curves) or larger (magenta curves) than median ages. For younger subjects, the distributions for the long (short) window ranges are peaking at ***α***_***DFA***_ values well above (below) the Gaussian expected value at ~0.5. For older subjects, however, the distributions were shifting significantly toward 0.5, indicating Gaussianity (one-sided Kolmogorov Smirnov, Bonferroni corrected, *p* < 0.05 for long windows; *p* < 0.01 for short windows; not significant for intermediate windows). Interestingly, the median ***α***_***DFA***_ *increased* with aging in the short windows range, correspondingly to reduced anti-persistence and *decreased* in the long windows range, reducing persistence. In other words, aging lead invariantly to replacing complexity with randomness.

Moving from group to subject-by-subject analyses, Fig. 5C shows the window size ranges for which correlations between subject-specific *α*_DFA_ and age were significant. For both empirical resting-state and task fMRI, correlations were significantly positive for short window sizes. There were also significantly negative correlations for several windows in the intermediate and long window size ranges. The range of significance was broader for resting-state than for task dFC streams.

We also performed a similar analysis for surrogate dFC streams. There were no correlations with age for shuffled dFC streams for which ***α***_***DFA***_ was always fluctuating around the Gaussianity value of ~0.5. For phase-randomized surrogates whose ***α***_***DFA***_-s were very close to the empirical data, although not precisely identical, we also found significant positive correlations in the short window range. However, the negative correlations with age in the intermediate and long window sizes ranges were tendential but not significant (even before applying multiple comparison correction). The replacement of complexity by randomness is thus more marked in empirical than in phase-randomized data.

Finally, in Fig. S7B we show that the ***α***_***DFA***_ computed from head motion time-series (cf. Fig. S7A) were not correlating significantly with age. Therefore, variations of ***α***_***DFA***_ with age for empirical dFC streams do not reflect –or at least, not uniquely– variations of head motion fluctuations (see *Discussion*).

### Dynamic Functional Connectivity correlates with task and cognitive performance

We tested whether dFC stream properties –as its speed ***V_dFC_*** and the scaling properties of its instantaneous increments summarized by *α*_DFA_– were indicative of behavioural or cognitive performance. We analysed in particular correlations with fluency in a simple visuomotor task or with the score obtained general in a common clinical assessment –the Monreal Cognitive Assessment (MOCA), probing several cognitive domains affected in age-related dementias (Nasreddine et al., 2005). In short, we found that speed of dFC measured during the task –but not at rest– correlated positively with task performance, measured by the spectral overlap (***SO***, Fig. 6A and Fig. S8A-B). We also found that *α*_DFA_ measured at rest correlated positively with the general composite MOCA score (Fig. 6B) and that variations of *α*_DFA_ between rest and task could correlate with visuomotor task PLI (Fig. S8C-D). More specifically, concerning dFC speed analyses, we found that window-range pooled task ***V_dFC_*** correlated with task’s ***SO*** for all three long (Fig. S8A), intermediate (Fig. 6A) and short (Fig. S8B) window ranges (bootstrap with replacement c.i., see captions for *p* values). Correlations were particularly strong in the intermediate window ranges. Considering single-window ***V_dFC,τ_***, we also found significant positive correlations over wide ranges of short, intermediate and long window sizes, as shown in Fig. 6C (blue ranges). Therefore, the faster were dFC streams during the task and the best manual movement could remain phase-locked with the displayed visual motion.

**Fig. 6.**
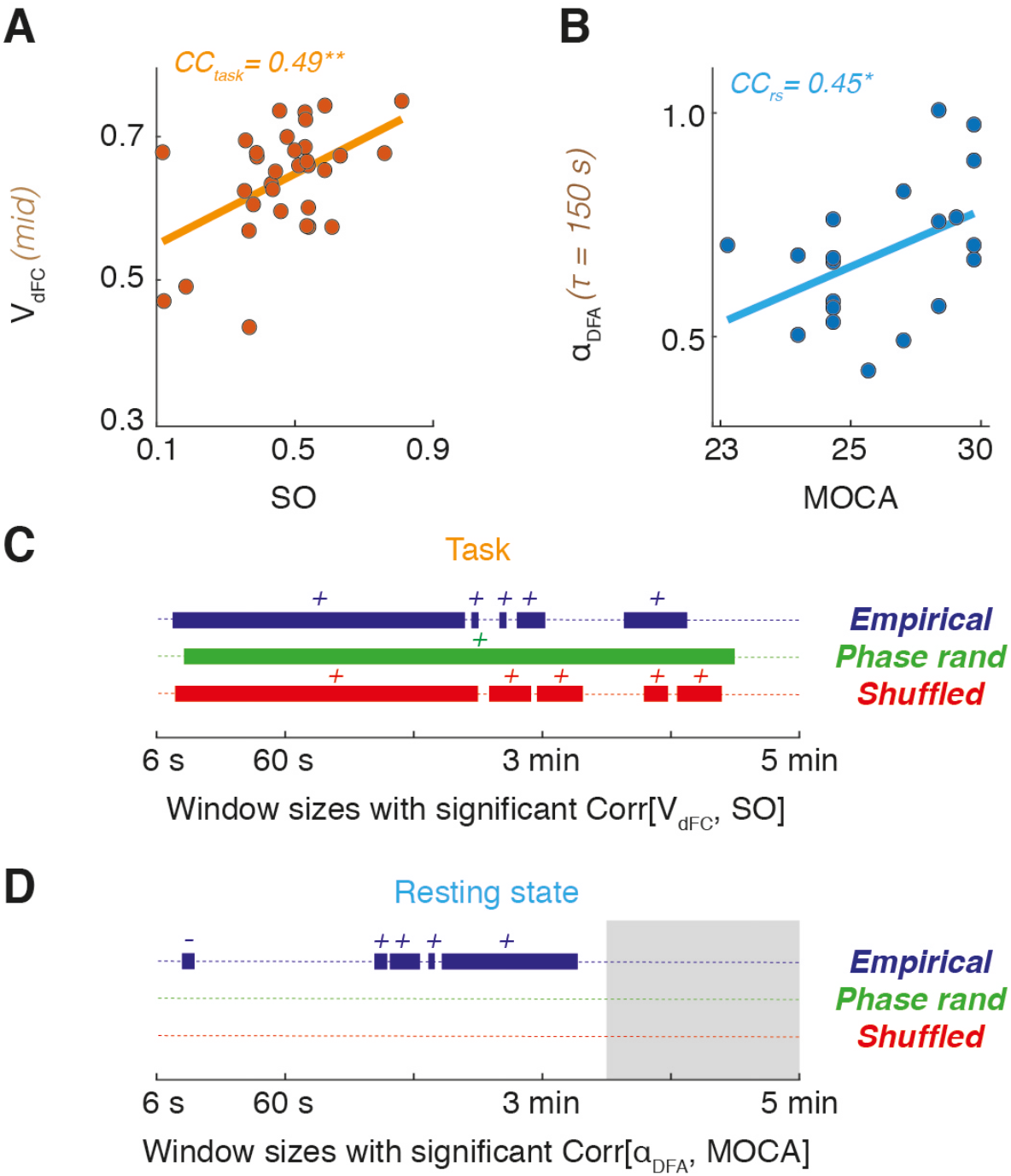
Correlation between dFC and cognitive/visuo-motor performance. Cognitive and behavioral efficiency were evaluated via a bimanual visuomotor task –in which a Spectral Overlap index (SO) was measured as a performance metric– and via a general cognitive assessment (summarized in a MOCA score). (A,B) Both SO and MOCA scores positively correlated with dFC metrics. In (A), we show a scatter plot of SO against the pooled dFC speed (here for empirical data in the intermediate speed range), measured during the task (bootstrap with replacement confidence intervals for Pearson correlation: **, *p* < 0.01). As shown by (C), significant (bootstrap, *p* < 0.5) correlations between SO and single window size dFC speed during task were found for broad window size ranges, not only for empirical data but also for both types of surrogates (a “+” sign indicates positive correlations). In (B), we show a scatter plot of the global MOCA score against α_DFA_ (here for empirical data at a representative window size in the long window sizes range), measured at rest (bootstrap with replacement confidence intervals for Pearson correlation: *, *p* < 0.05). As shown by (D), significant (bootstrap, *p* < 0.5) correlations between resting-state α_DFA_ and MOCA score are found only for empirical data and not for surrogates, in selected window size ranges (a “-” or “+” sign indicate negative or positive correlations). The max τ used for single-window α_DFA_ calculation was shorter than for dFC speed analyses and trimmed to 210 s (excluded τ range shaded in gray)

The correlations between dFC speed and ***SO*** also held for surrogate task dFC streams of both phase-randomized and shuffled types, over window size ranges close to the ones for empirical data (Fig. 6C, red and green ranges). Therefore, as in the case of correlations with age (Fig. 4D), it seems that ***SO*** correlations with dFC speed can be accounted for by a general increase of dFC variance, which is preserved when constructing the surrogates dFC streams (see *Discussion*). Correlations between resting-state dFC speeds and ***SO*** were not significant.

Concerning DFA analyses, we found (Fig. 6D) that resting-state *α*_DFA_ correlated positively with the general MOCA score over a substantial range of window sizes located within the long window size range (τ > 90 s). There was also a small range within the short windows range for which ***α***_***DFA***_ was negatively correlated with MOCA. Therefore, the more the dFC stream deviated from memoryless random walks and the higher was the composite MOCA score. Remarkably and differently from dFC speed analyses, we found that these correlations between ***α***_***DFA***_ and MOCA reached significance only for empirical data but not anymore for surrogate data, including phase-randomized data. Note also that the MOCA scores were quite diverse across the subjects and that the correlation between age and MOCA was mildly negative but not significant. Therefore, correlations between subject-specific ***α***_***DFA***_ and MOCAs seem to depend on fine details of the empirical dFC increment sequence rather than on generic statistical aspects rendered as well by the phase-randomized surrogates.

The task ***α***_***DFA***_-s were generally different from the rest ones, sometimes larger and sometimes smaller. It was thus possible to calculate for each subject the quantity **Δ*α***_***DFA***_, positive or negative depending on the subject, that could be negative or positive depending on the subject. In Fig. S8C, we show a scatter plot of task-vs-rest **Δ*α***_***DFA***_ against ***SO*** for a representative window size. The more the scaling exponent increases –get more persistent-during tasks relative to rest and the better the visuomotor task will be performed. In Figure S8D we show the windows for which positive correlations between **Δ*α***_DFA_ and ***SO*** remained significant. This range, once again located within the long window sizes range, is smaller than for correlations between ***α***_***DFA***_ and MOCA. However, these correlations only appeared in the empirical data but not in the two surrogate streams.

## Discussion

We characterise dFC by allowing ***FC(t)*** fluctuations to be interpreted as a stochastic walk in the space of possible network configurations. By this, dFC forms a continuous stream without the need to segment epochs belonging to sharply separated “FC states” or clusters. We focused on two aspects: the speed at which the stochastic exploration of the FC space is performed and the geometry of the resulting stochastic walk.

As for speed, the key idea is that variability of ***FC(t)*** occurs at all times and not necessarily just restricted to specific switching events. In this sense, our approach quantifying rate of continuous variation along the dFC stream is conceptually akin to temporal derivative methods introduced by Shine et al. (2015) or used in EEG analyses (Schurger et al., 2015; He, 2018). However, these previous studies addressed variations at the level of signals or activation states, rather than at the FC network level. Distributions of dFC speed were clearly unimodal for all considered subjects, through all the probed window sizes and for both task and rest. The peak of the dFC speed distribution could thus be interpreted as a “typical dFC speed”, a first indicator of the expected scale of time-to-time network variability along the dFC stream. We found that this typical speed decreases with age, and this not only for empirical rest and task data but also for both considered surrogate types. Overall, our findings hint toward a reduced network variability in aging, in the same direction as other dFC analyses (Chen et al., 2017) and previous reports of reduced variability in elderly already at the level of the BOLD signal itself (Grady & Garrett 2014). Other studies, however, reported increased “noise” in the elderly relative to the younger subjects, at least at specific scales and in certain regions (Yang et al., 2013). In even stronger apparent conflict with our findings are studies indicating that, along dFC, network nodes tend to fluctuate between network modules more dynamically in elderly than in younger subjects, resulting in a more flexible modular structure (Schlesinger et al., 2016; Davison et al., 2016). Such divergences are not necessarily in contradiction with our results. Our metric of dFC speed is a mere correlation distance at the level of whole network comparison. First, being normalized, it is not sensitive to variations in signal variances, as long as the signal correlation structure is preserved. Second, enhanced flexibility of modules also indicates that, overall, the separation between modules observed in young is blurred by node exchange, resulting in decreased within-module and increased between module average connectivity (Betzel et al., 2014). Thus, the overall differences between time-resolved network frames with soft modules (with different “nuances of gray”), as observed in the elderly, could be smaller than the differences between frames with neat modules (“black or white”), as observed in the young.

Even if network reconfiguration never stops, the rate of reconfiguration is not constant. Analyses focusing on the detection of non-stationarity and state changes in FC, emphasized the role of long jumps –which we called here dFC leaps– across which the variance is so large to be interpreted as a significant change of the FC network link strengths (see e.g. Zalesky et al., 2014; 2015). Here, we instead insist on the fact that short jumps are abundant in empirical resting-state and task data relative to surrogate dFC streams in which the shuffling of timeframes removes sequential correlations. The “agglutination” of these many small variation events results in temporally connected epochs of transient slowing down. Famously, even the tortoise can beat the fast runner Achilles. Through the concatenation of many short steps, it becomes possible, indeed, to travel overall long distances. Thus, dFC knots are not necessarily indicating the existence of compact clusters separated by long gaps (cf. the comparison between Figs. 1C and 1D). Still, it may be that at least some dFC leaps are large enough to be qualified as proper FC state transitions (cf., the over-expression of large speeds in Fig. 3). We insist here once again on the fact that the debate “stationary vs. non-stationary” (see e.g., Hindriks et al., 2016) may not be the more pertinent when attempting to capture the nature of dFC specificities. It instead appears that what makes empirical dFC unique relative to surrogate ensembles are its specific sequential correlation properties, deviating from trivial Gaussian stationarity.

We used fractal scaling analysis to assess “how” empirical dFC streams manifest long-range sequential correlations. We identified for most window sizes deviations of empirical dFC streams from Gaussianity, finding strong evidence for long-range correlations, positive –denoting persistence– or negative –denoting anti-persistence– depending on the window size. We detected a nearly identical window-size dependent spectrum of long-range correlations for a fully independent dataset mediated from the HCP initiative, hinting at the fact that our findings are general for resting state fluctuations of the BOLD signal and not just restricted to our specific study.

A possibility is that higher than Gaussian ***α***_***DFA***_ exponents are the ultra-slow range extension or at least the surviving observable shadow of faster scale-free microstate fluctuation dynamics (Van de Ville et al., 2010) that cannot be tracked with the too-low temporal resolution of fMRI. We observed this persistence of dFC streams very robustly over the whole intermediate and long window size ranges. Over time windows from ~20s up to several minutes, exponents are safely lying above the Gaussian value of ***α***_***DFA***_~0.5. The extension of the range for which persistence holds also means that, although the exact dFC speed and ***α***_***DFA***_ values may quantitatively change for different window sizes, our random walk analyses are describing essentially the same qualitative phenomenon, as long as the adopted window size is larger than ***τ*** > 20s.

Long-range persistence has been observed since long in contexts as diverse as water flooding (Hurst, 1951; Mandelbrot & van Ness, 1968), fluctuations of musical rhythms (Hennig et al., 2011), foraging in ecosystems (Viswanathan et al., 1999), prices on the stock market (Lo, 1991), internet traffic (Cleveland and Sun, 2000), etc. In neuroscience; long-term persistence has been identified in resting-state and task fluctuations of electrophysiological and imaging signals (Linkenkaer-Hansen et al., 2001; Van de Ville et al., 2010; He, 2014) and found to correlate with behaviour (Palva et al., 2013; Ciuciu et al., 2014). Anti-persistence is less frequently discussed, but it has also been routinely found, for instance, once again in nonlinear dynamics (Penna et al., 1995), sportive scores series (Gabel and Redner, 2011) or in heartbeat (Peng et al., 1993), more relevant here since it may be a potential contaminating artefact of physiological but not neural origin in the BOLD signal.

As a matter of fact, our observation of multifractal scaling in dFC is not completely unexpected. Quite on the contrary, it is well known that fMRI time-series at rest and during tasks display very characteristic multifractal spectral properties, relating to both behavioural performance and pathological alterations and modulated by task difficulty or aging (Maxim et al., 2005; Ciuciu et al., 2012; He, 2014; Churchill et al., 2016; Dong et al., 2018). Tools way more sophisticated than the ones we adopt here have been used to characterize and confirm multifractality in fMRI signals (Ciuciu et al., 2017; La Rocca et al., 2018). Furthermore, the multifractal properties of fMRI signals hold not only at the level of univariate spectra but also of cross-spectra therefore translating into specific signatures even at the level of static networks, once again with behavioural correlates (Ciuciu et al., 2014). Therefore, our findings reconnect with solid, cumulating evidence about the fact that fluctuations of coordinated neural activity have non-Gaussian components. Most likely, the multifractality of the original underlying signals contributes to the multifractality of the derived dFC streams. It is however only when discussing multifractality directly at the higher level of time-resolved networks, that the interpretation of dFC as an anomalous random walk in FC space naturally arises. Beyond merely proposing variants of previously used quantitative biomarkers, the fact itself of lifting up multifractal analyses from the level of activations to the level of the “chronnectome” enables a qualitatively new vision of what dFC really is: a non-random, complex search for a coordinated brain state –captured by the transient FC*(t)*– within a large dimensional space of possible dynamic configurations.

Persistent random walks have been associated to optimality in local search for resources during foraging in ecosystems (Viswanathan et al., 1999) but also for “mental space search” of optimal strategies during bidding (Radicchi et al., 2012). Intriguingly, we observed that higher *α*_DFA_ exponents –i.e. deviating more from unstructured Gaussian random walk and more approaching persistent walks of the Levy type (Metzler et al., 2014)– were associated with better general cognitive performance (Fig. 6B and 6D). Following up on our interpretation of dFC as an anomalous stochastic walk, we may thus speculate that dFC serves as a neural process to efficiently ‘forage’ for cognitive processing resources. Specifically, the non-trivial fluctuations of FC*(t)* may play the functional role of searching for FC patterns adapted to the information routing demands (Battaglia et al., 2012; Kirst et al., 2016) of ongoing mental computations.

The anti-persistence of dFC at very short window-sizes also correlated with cognitive performance. However, this correlation was not anymore holding at the partial correlation level when we regressed age out as a common covariate (unlike the stronger positive correlations found for intermediate and long window ranges, which robustly survived even at the partial correlation level). This anti-persistence was not specific to empirical dFC, since it was found as well in phase-randomized surrogates and varied with age. Since “blue” anti-persistent noise is associated to heart-beat variability (Peng et al., 1993) and that the fractal scaling of heart-beat dynamics is also affected by aging (Iyengar et al., 1996) –as well as in the case of many other physiological signals (Goldberger et al., 2002)– is likely that dFC estimation over these very short scales is affected by physiological artefacts of not neural origin. However, persistence observed for window size ***τ*** > 20 s is more robust, and the measured DFA exponents are very different from the ones concomitantly measured from head-motion (cf. Fig. S7).

Ultimately, a strong indication of the non-artefactual nature of empirical dFC is the fact that its random walk properties do correlate to a certain extent with behaviour and cognitive performance. Theories of cognitive aging have advanced that a cause for declining performance would be the insufficient access to cognitive resources due to a reduced speed of information processing (Salthouse, 1996; Finkel et al., 2007). Cognitive aging has been associated with deficits in disengaging from active brain functional states, more than to alterations of the states themselves (Clapp et al., 2011; Cashdollar et al., 2013). Aging affects dFC streams by reducing their speed. Thus, it may seem that slowing down of cognition is paralleled by slowing of dFC. To move beyond mere conjectures, we do believe that more and better-adapted experiments should be designed to probe speed of processing or task switching. Indeed, the fact that dFC speed correlations with task performance are significant during task fMRI but only tendential during resting-state may also hint at the fact that dFC “moves slower” because motion during the task is different, if not slower. Theoretical studies on task dynamics in behavior and their neural correlates emphasize the emergence of low-dimensional subspaces holding task-specific flows (Kelso 1995; see Huys et al. 2014 for a review), which act as generative models for cognitive and behavioral processes. Structured Flows on Manifolds (SFMs) are the mathematical representation of the deterministic features underlying behavior, including multistability, convergence/divergence of trajectories, as well as task-specific stability and robustness (Pillai et al. 2017). They arise from interactions of coordinated brain activations within the neuro-skeletomuscular system including visuomotor tasks (Fink et al. 2008; Sleimen-Malkoun et al., 2014), multi-limb coordination (Kelso 1995; Fink et al. 2000a, 2000b), multisensory integration (Lagarde et al. 2006) and learning (Schöner et al. 1992; Zanone et al. 1997), but causal contributions are difficult to disentangle. It is tempting to interpret the “randomness replacing complexity” pattern observed here at the level of DFA analyses as degradations of SFMS and a novel form of de-differentiation, a collective name encompassing several kinds of destructuring and loss of complexity occurring in healthy aging at the level of behavior, cognitive strategies and brain activity (Baltes et al., 1980; Lemaire & Arnaud, 2008; Sleimen-Malkoun, 2014).

Interestingly, we also found that performance in our visuomotor task correlated with the capacity to actively modify the DFA exponent found at rest toward a different functioning value during task (Fig. S8C-D). Our finding suggests that brain systems actively tailor the scaling properties of emergent dFC to the specific task demands and that the capacity to do so is important to explain the achieved performance. Similar results have been found at the level of changes in the scaling properties of static FC correlations (Ciuciu et al., 2014), and we now further extend them at the level of dFC analyses. With our dataset and analyses did not allow for probing these and other hypotheses beyond speculations. In particular, the MOCA global cognitive score provides only a very rough summary characterization of the many facets of cognitive performance and its decline. To probe relation between random walk features of dFC and cognitive performance, one would need a more systematic and controlled cognitive testing setup, probing a spectrum of different cognitive functions (attention, working memory, their executive control, etc.) under different conditions within the same subject. Furthermore, it is unlikely that cognitive performance levels in specific tasks –beyond generic global assessment of performance– is associated to alterations of dFC properties at the whole brain level. Indeed, different specific tasks differentially involve alternative functional sub-networks.

To this heterogeneity of regional involvement, may correspond a parallel heterogeneity of dFC properties, which we fully ignore in this first study. It is known that variance of FC links in time (Chen et al., 2017) or even the fractal scaling properties of fMRI signals (Maxim et al., 2005; Churchill et al., 2016; Dong et al., 2018) are affected heterogeneously across brain regions. In a companion study by Lombardo et al. (2020) we make a further step forward, probing variations of cognitive performance in selective attention and other cognitive functions, before and after sleep deprivation. We do so by developing “modular” (i.e. subnetwork specific) forms of dFC speed analysis. We can thus show that, beyond the first –useful but rough– approximation of whole-brain dFC analyses, distinct subsets of functional links can evolve with different dFC speeds. Furthermore, differential modulations of modular speed can result in differential modulation of performance through different tasks. A key result of Lombardo et al. (2020) is that, when cognitive performance changes are small and more task-specific than the large overall cognitive performance losses linked to MOCA score decreases, then they cannot be captured anymore by whole-brain dFC analyses. They can however still be tracked by modular dFC analyses. The reader is referred to Lombardo et al. (2020) for more details.

Our random walk analyses may be combined with improved ways of estimating dFC streams themselves, for instance, by avoiding the use of sliding windows (Lindquist et al., 2014; Yaesoubi et al., 2018). More importantly, it would be important to investigate potential mechanisms giving rise to complex temporal structure between order and disorder in empirical dFC streams. The emergence of long-range correlations has often been associated with critical dynamics (Linkenkaer-Hansen et al., 2001; Chialvo, 2010), and resting-state is well captured by dynamic mean-field models close to a critical point (Deco et al., 2011). Various proposals have been made for the potential clinical use of dFC (McIntosh et al. 2019) and related metrics (such as DFA and multiscale entropy) quantifying the changes of flow and manifold structure across pathological conditions. More in general, long-range correlations are also generated by non-linear behavior at the “edge of chaos” (Manneville, 1980; Geisel et al., 1987), well in line with our observation that the resulting dFC lies between order and randomness, as also observed in spontaneous activity at the micro-scale (Clawson et al., 2019). Mean-field whole-brain computational models, besides providing further confirmations of the non-artifactual nature of dFC – virtual brains do not have blood –, may allow identifying the dynamic and neurophysiological mechanisms behind its random walk properties. Generic whole-brain models are already able to qualitatively reproduce switching dFC (Hansen et al., 2015; Cabral et al. 2017a), but enhanced dynamical complexity will be required to account *in silico* for the rich non-linearities of empirical dFC revealed by our approach.

Future simulations might be fitted to individual subjects via automated pipelines (Schirner et al., 2015; Proix et al., 2016) to render the dFC trajectory of evolution across aging more quantitatively. Models embedding SC typical of different age classes may reproduce the slowing down and complexity loss of dFC as an emergent by-product of SC ‘disconnection’ itself (Salat, 2011). Or, more likely, they may show that this disconnection must be compensated to account for observations by a drift of the global ‘dynamic working point’ of operation of cortical networks, which could be possibly induced by altered neuromodulation (Bäckman et al., 2006) or metabolism (Arenaza-Urquijo et al., 2013).

## Abbreviations

*rs*: resting-state
*RSN*: resting-state network
*fMRI*: functional magnetic resonance imaging
*BOLD*: blood oxygen level dependent
*SC*: structural connectivity
*FC*: functional connectivity
*d*FC**: dynamic functional connectivity
*DFA*: detrended fluctuation analysis
*MoCA*: Montreal Cognitive Assessment
*SO*: spectral overlap

## Author contributions

DB and VJ conceived the novel analytic methods; DB and TB performed imaging data analysis; DL and EH contributed to dFC toolbox development; AD provided expertise and toolbox for DFA and model selection; PR supervised imaging experiments; PR, AD and SC conceived the cognitive and motor experiments; SC performed the cognitive and motor experiments and processed behavioral data; RM and JZ contributed to data pre-processing; DB, TB, DL, EH, SC, AD, RM, JZ, PR and VJ wrote the paper.

## Acknowledgments

This research was supported by the Brain Network Recovery Group (through the James S. McDonnell Foundation) and by the European Union’s Horizon 2020 Framework Program for Research and Innovation under the Specific Grant Agreement No. 785907 (Human Brain Project SGA2). DB acknowledges support from the Mission for Interdisciplinarity of the CNRS (Infiniti program 2017-2018, “BrainTime”), from the EU Innovative Training Network “i-CONN” (H2020 ITN 859937). DL has been funded by the Uruguayan National Agency of Research and Innovation (ANII, grant POS EXT 2015 1 123495). P.R. acknowledges the following additional funding sources: H2020 Research and Innovation Action grants VirtualBrainCloud 826421 and ERC 683049; German Research Foundation CRC 1315, CRC 936 and RI 2073/6-1; Berlin Institute of Health & Foundation Charité, Johanna Quandt Excellence Initiative. We thank: Rodrigo Sigala, Sebastian Haufe, Michael Schirner, Simon Rothmeier for help in data acquisition; and Djouya Arbabyazd, Dionysios Perdikis, Rita Sleimen-Malkoun, Annette Witt and Paul Triebkorn for discussions.

## Supplementary material

### Supplementary figures

**Fig. S1.**
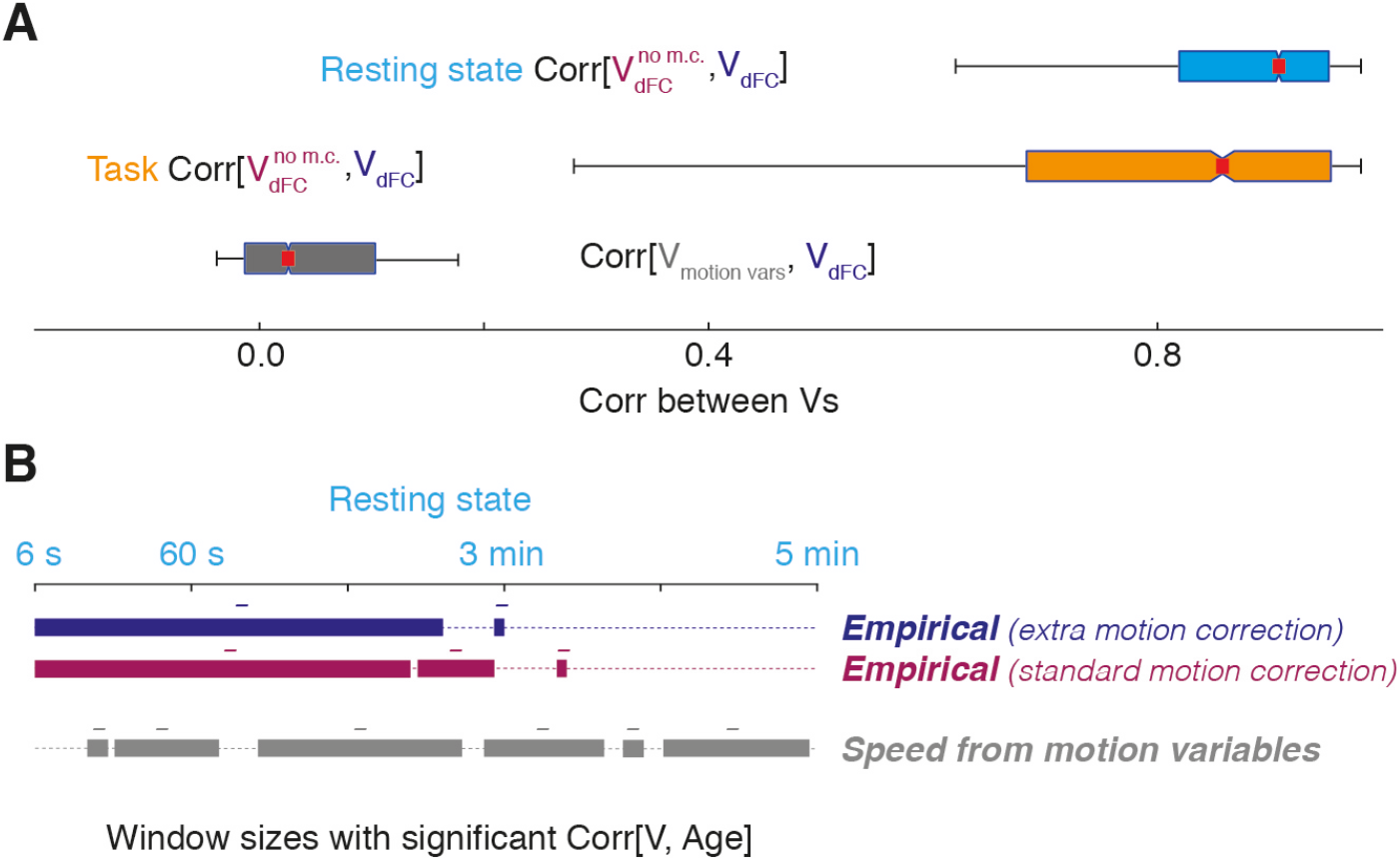
dFC speed analyses are robust against motion artifacts. To verify the potential contamination of our dFC speed estimations by motion artifacts we studied the statistics of fluctuations of head displacement variables themselves as a function of age and correlated them to dFC speed fluctuations, evaluated with two alternative motion correction strategies: a weaker one in which a standard motion correction pipeline is adopted (providing speed estimations denoted here as 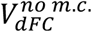); and the stronger one used in the main article (providing speed estimations denoted as *V_dFC_*) in which, additionally, head motion variables are explicitly regressed out the signal after correcting it with standard measures. (A) Correlations between dFC speeds estimated with the weaker or the stronger motion correction strategies generally remain high for both resting-state (light blue) and task (orange). On the contrary, estimating “speed” directly on the stream of 6-variate head motion variables to track the alternation of smaller and larger head excursions, lead to motion dFC-speed-like sequences which are only poorly correlated with properly motion-corrected dFC speed *V_dFC_* (grey). Therefore, dFC dynamics do not reflect head motion dynamics. Boxplots are constructed for speed-to-speed correlations for all window-sizes pooled. Boxes denote the interquartile range, whiskers the range between the 5% and the 95% sample percentiles. A red mark indicates the median, surrounded by a notch, which reflects Kruskal-Wallis testing of medians, significantly different (*p* < 0.05) if notches are not overlapping. (B) There are significant negative correlations (bootstrap, *p* < 0.05) between age and dFC-speed-like quantities estimated directly on motion variables, i.e. median head displacement speed decreases with age. However, the ranges of window size in which age correlations hold for head motion dFC-speed-like quantities and proper dFC speeds, with both types of motion correction protocols, are very different, suggesting that age effects captured at the level of dFC streams are not merely age variability of head motion.

**Fig. S2.**
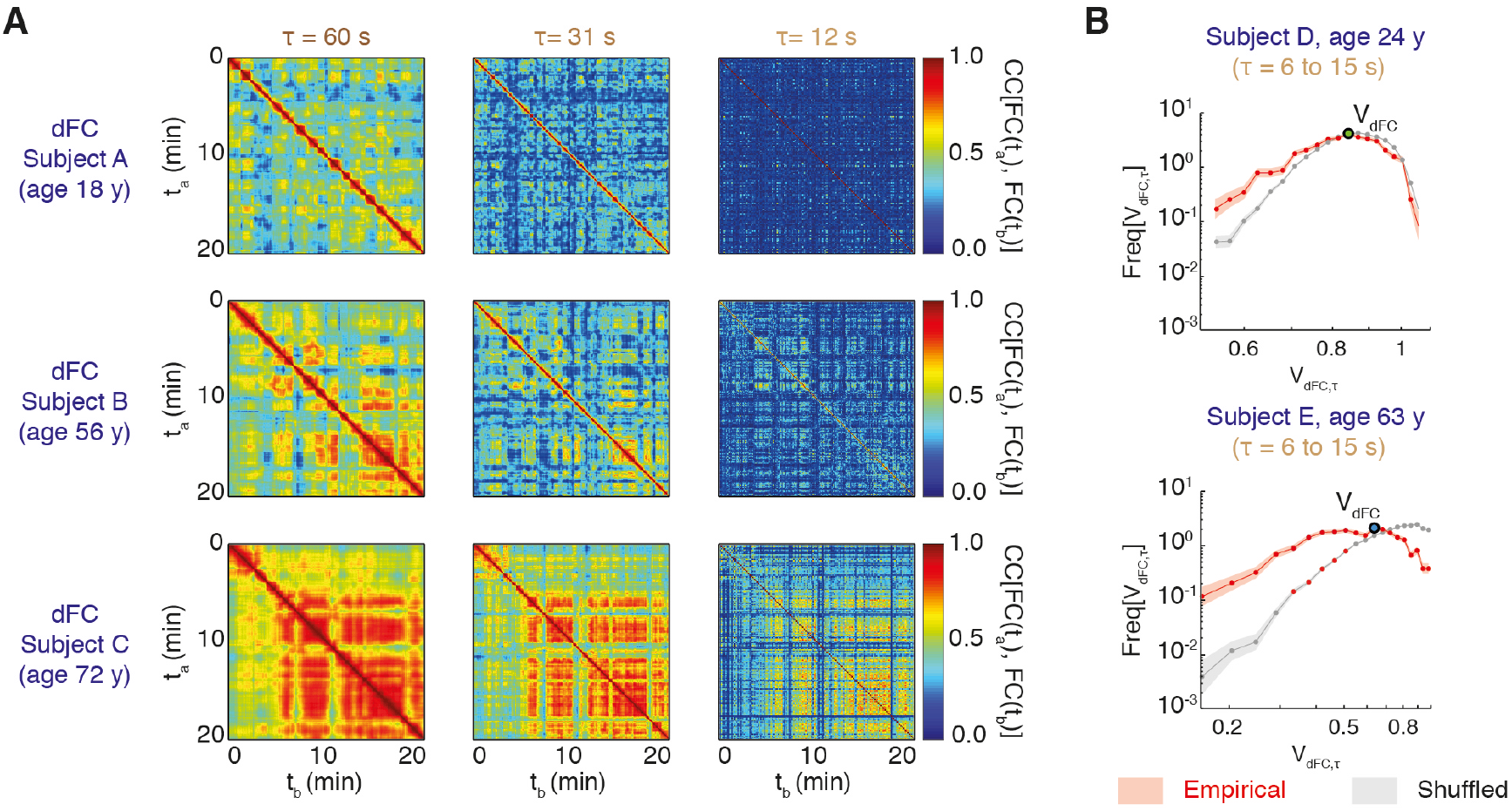
dFC matrices for different sliding window sizes and single subject dFC speed distributions. (A) dFC analyses for the same subjects considered in Fig. 4A are shown, for three different dFC window sizes τ-s. To simplify comparison, in the left column we reproduced the same dFC matrices already presented in Fig. 4A. Blocks of relatively elevated inter-network correlation corresponding to dFC knots were visible for all time scales and ages. (B) Singe-subject distributions of dFC speed, shown here for two representative subjects (log-log scale, pooled window sizes 6 s ≤ *τ* < 15 s, corresponding to the short windows range) displayed a peak at a value *V_dFC_* (*typical dFC speed*) and a fat left tail, reflecting an increased probability relative to chance level (shuffled surrogate, null hypothesis of “randomness”) to observe short dFC steps (95% confidence intervals are shaded: red, empirical; gray, chance level from shuffled surrogates).

**Fig. S3.**
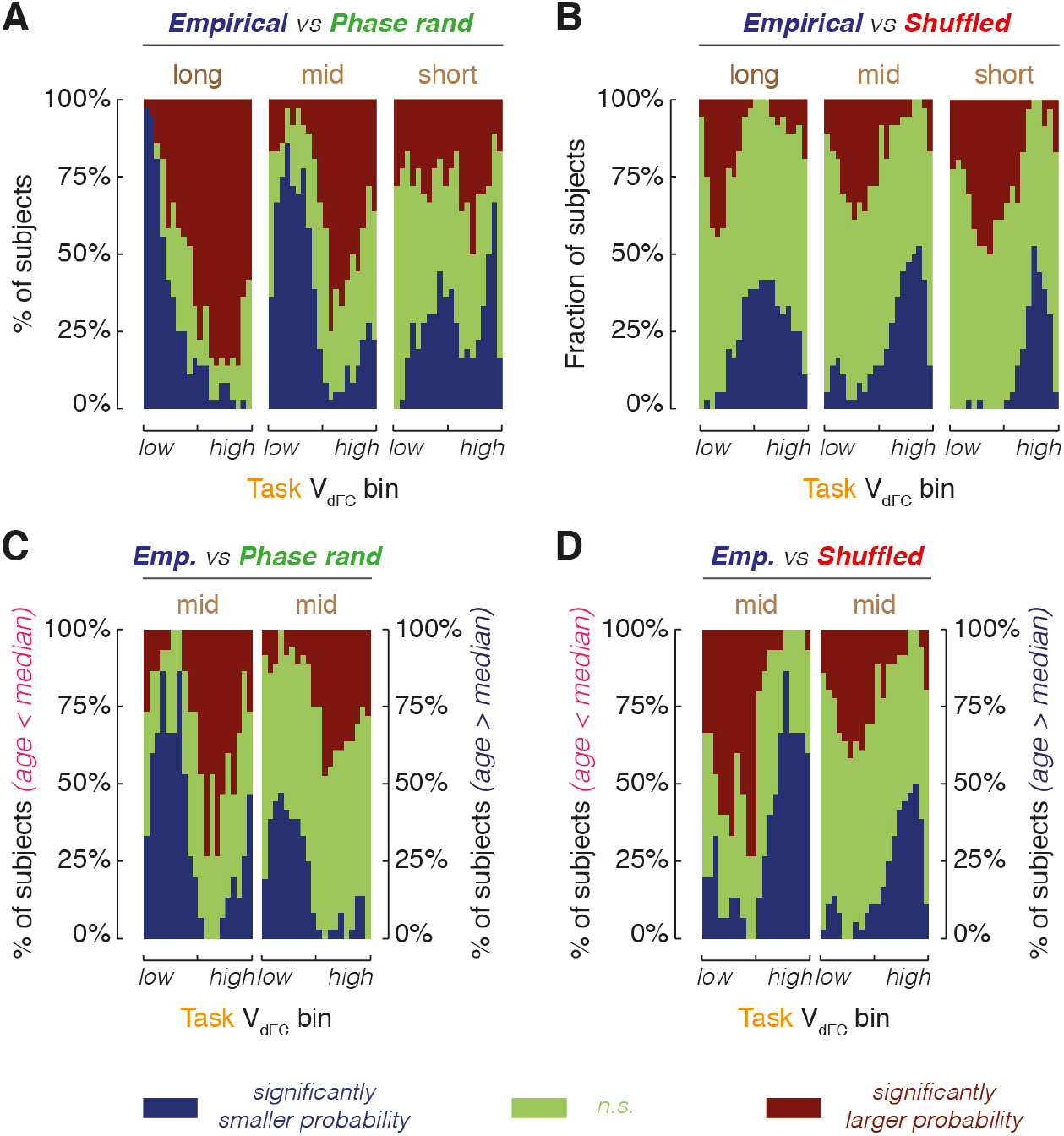
Empirical dFC streams lie “between order and randomness” *(task analyses)*. Analyses identical in all parameters to Figure 3 but based on single subject dFC speed distributions sampled during task blocks. All results are confirmed. (A) Comparison with phase-randomized surrogates indicates that, in empirical task data, lower (higher) than median dFC speeds are often under-(over-) represented. These effects are particularly evident in the long time-windows range (leftmost plot). (B) Comparison with time-shuffled surrogates indicates that, in empirical task data, lower (higher) than median dFC speeds are often over-(under-) represented, i.e. a reverse pattern with respect to phase-randomized surrogates. (C-D) When separating subjects into two age groups (younger or older than the median), the comparison patterns revealed by panels A and B are confirmed, but crisper for young subjects and more blurred for older subjects. Overall, if we dub as “order” the null hypothesis of static average FC (i.e. phase-randomization) and as “randomness” the null hypothesis of temporally uncorrelated dFC fluctuations (i.e. time shuffling), the statistics of task dFC fluctuations appear to lie “between order and randomness” (i.e., they are “complex”).

**Fig. S4.**
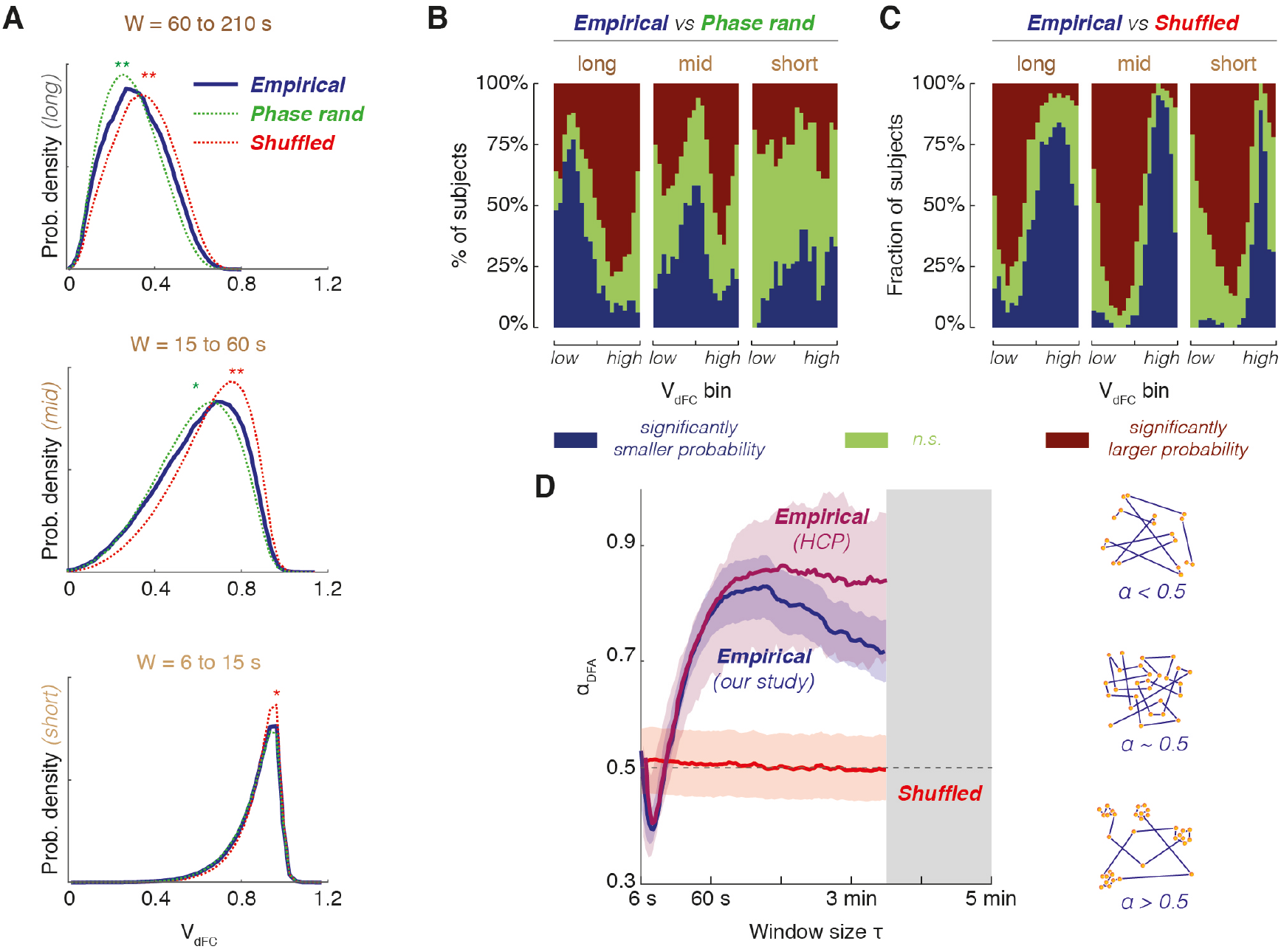
Empirical dFC streams lie “between order and randomness” *(control resting state dataset)*. Analyses identical to Figs. 2G, 3A-B and 5A but performed on a control resting state fMRI data mediated from the Human Connectome Project for comparison with our original aging dataset. (A) Distributions (smoothed kernel-density estimator) of resting-state dFC speeds for empirical and surrogate ensembles (averaged over subjects), pooled over three distinct window-size ranges (statistical testing, significance conventions and window-size ranges as in Fig. 2G). (B-C) Comparisons of empirical and surrogate histograms of resting-state dFC speeds at the single-subject level. As in Figs. 3A and 3B, we perform comparisons speed bin by speed bin, checking whether dFC speeds within a given bin are observed with a probability significantly above chance-level (red), significantly below chance level (blue) or compatibly with chance level (green), according to a stationarity (B) or lack of sequential correlations (C) null hypotheses. (D) Detrended Fluctuation Analysis (DFA) of the time-series of instantaneous variations of FC*(t)* along resting-state dFC streams. For every window size, we quantify the DFA exponent α_DFA_ indicating whether dFC fluctuations have long-range persistence (0.5 < α_DFA_ < 1), long-range antipersistence (α_DFA_ < 0.5) or are compatible with a Gaussian random walk (α_DFA_ ~ 0.5). Lines report the median α_DFA_ (purple color for the HCP control dataset and, for comparison, our aging dataset in blue) and shaded contours the 95% confidence interval over subjects (median ± 1.96*standard deviation of sample mean). Distributions of dFC speeds, at the group or single subject levels, as well as DFA exponents for this HCP control dataset are qualitatively equivalent to our aging dataset.

**Fig. S5.**
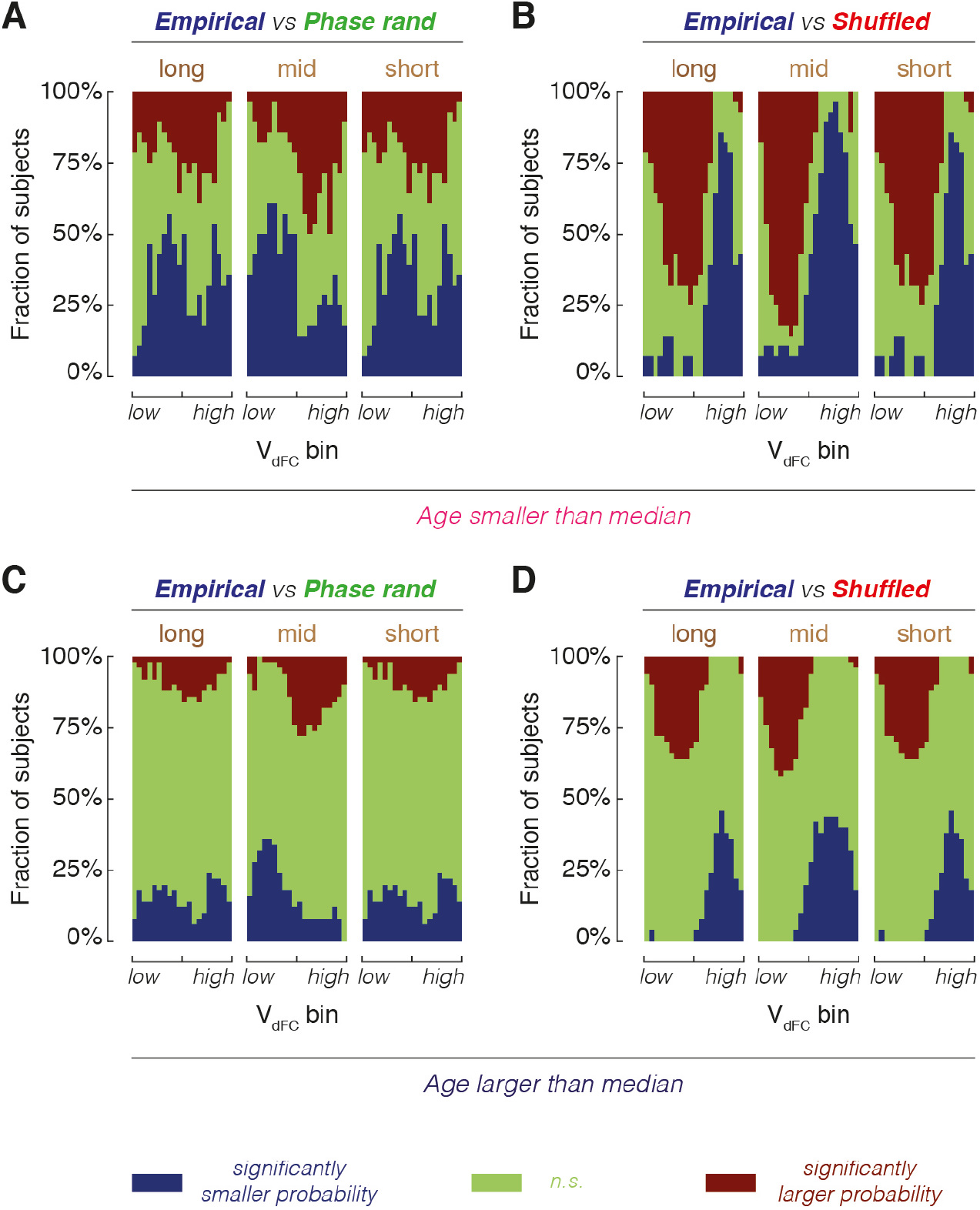
Empirical dFC streams lie “between order and randomness” *(additional resting-state analyses)*. As in Figure 3, we compare here empirical and surrogate histograms of resting-state dFC speeds, pooled over three different window-size ranges (long, intermediate and short) at the singlesubject level. The plots shown here are constructed as the ones presented in Figure 3 but age-separated analyses (A-B, for “young” group; C-D, for “older” group) are shown for all the three pooled window ranges (long to short, from left to right subpanels), for comparisons with both types of surrogate (empirical vs. phase-randomized in panels A and C; vs. time-shuffled in panels B and D). The deviation of empirical dFC speed distributions from the “ordered” phase-randomized surrogates and the “random” time-shuffled surrogates get more blurred with aging across all probed window size ranges.

**Fig. S6.**
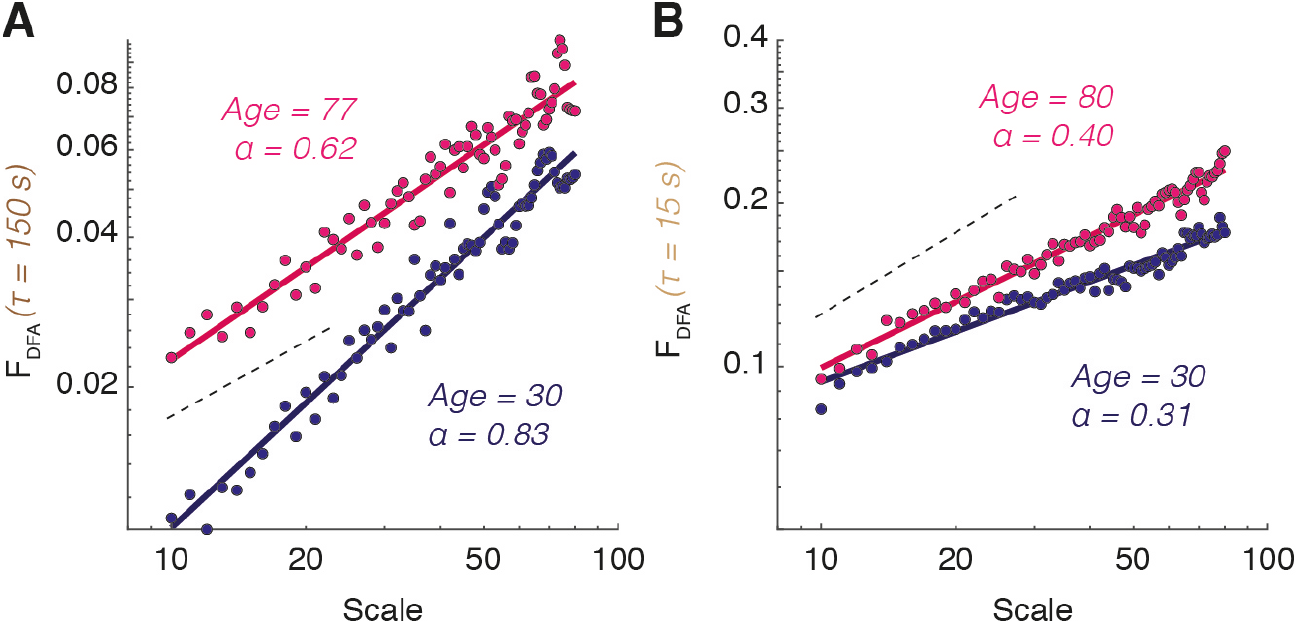
Genuine power-law scaling in DFA analysis. Shown here are examples of DFA log-log plots for representative subjects (in the “young” group, in blue; or in the “older” group, in magenta) and two representative window sizes for dFC stream estimation (a τ in the long window sizes range for panel A and a τ in the short window sizes range for panel B). The linearity of DFA scatter plots on the log-log plane (scale of coarse-graining vs total fluctuation strength), confirmed by Bayesian model comparison, reveals that instantaneous increments along the dFC stream form a self-similar sequence. The black dashed lines indicate the slope that would be associated to α_DFA_ = 0.5, i.e. the case of ordinary uncorrelated Gaussian random walk (A) For the shown DFA plots, linear slopes are steeper than for ordinary random walk, indicating that dFC streams evaluated at this long window size follow a persistent stochastic walk. (B) For the shown DFA plots, linear slopes are less steep than for ordinary random walk, indicating that dFC streams evaluated at this short window size follow an anti-persistent stochastic walk. In both panels A-B, linear slopes for the older subjects are closer to an ordinary random walk (“randomness” replaces “complexity” with aging).

**Fig S7.**
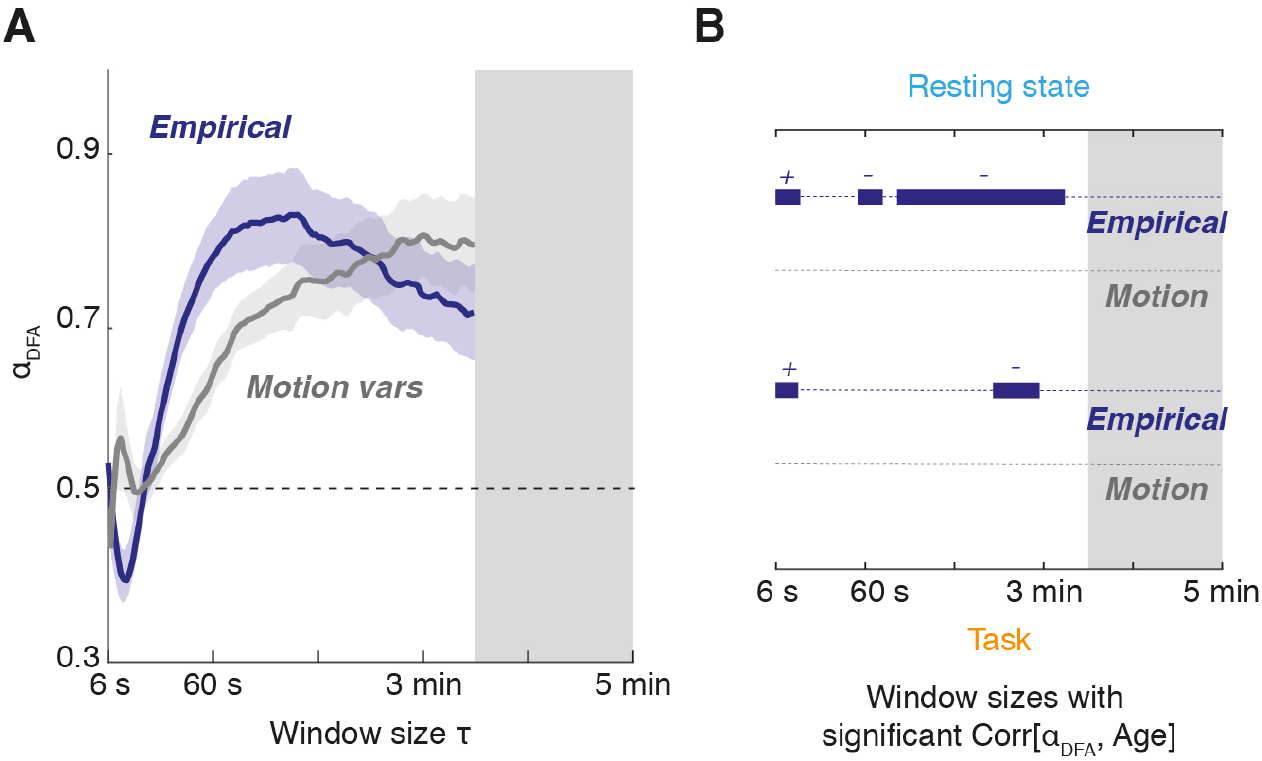
DFA analyses are robust against motion artifacts. (A) We performed Detrended Fluctuation Analysis (DFA) of the multivariate time-series of head motion (grey color). Reported is also the analogous line for resting-state empirical fMRI data, copied from Figure 5A. Lines report the median α_DFA_ and shaded contours the 95% confidence interval over subjects (median ± 1.96*standard deviation of sample mean). The max τ used for single window α_DFA_ calculation was shorter than for dFC speed analyses and trimmed to 210 s (excluded τ range shaded in gray). The spectrum of α_DFA_ exponents for motion time-series is completely different from BOLD fMRI, although is also deviating from normality. (B) The DFA exponent for motion variables never correlates significantly with age at any of the tested time-windows. We report for comparison as well the ranges in which fMRI BOLD α_DFA_ exponents correlate significantly with age, copied from Figure 5C (a “+” or “-” sign indicate respectively positive or negative correlation). Above, resting state; below, task dFC speeds (significance tested via bootstrap with replacement, *p* < 0.05).

**Fig. S8.**
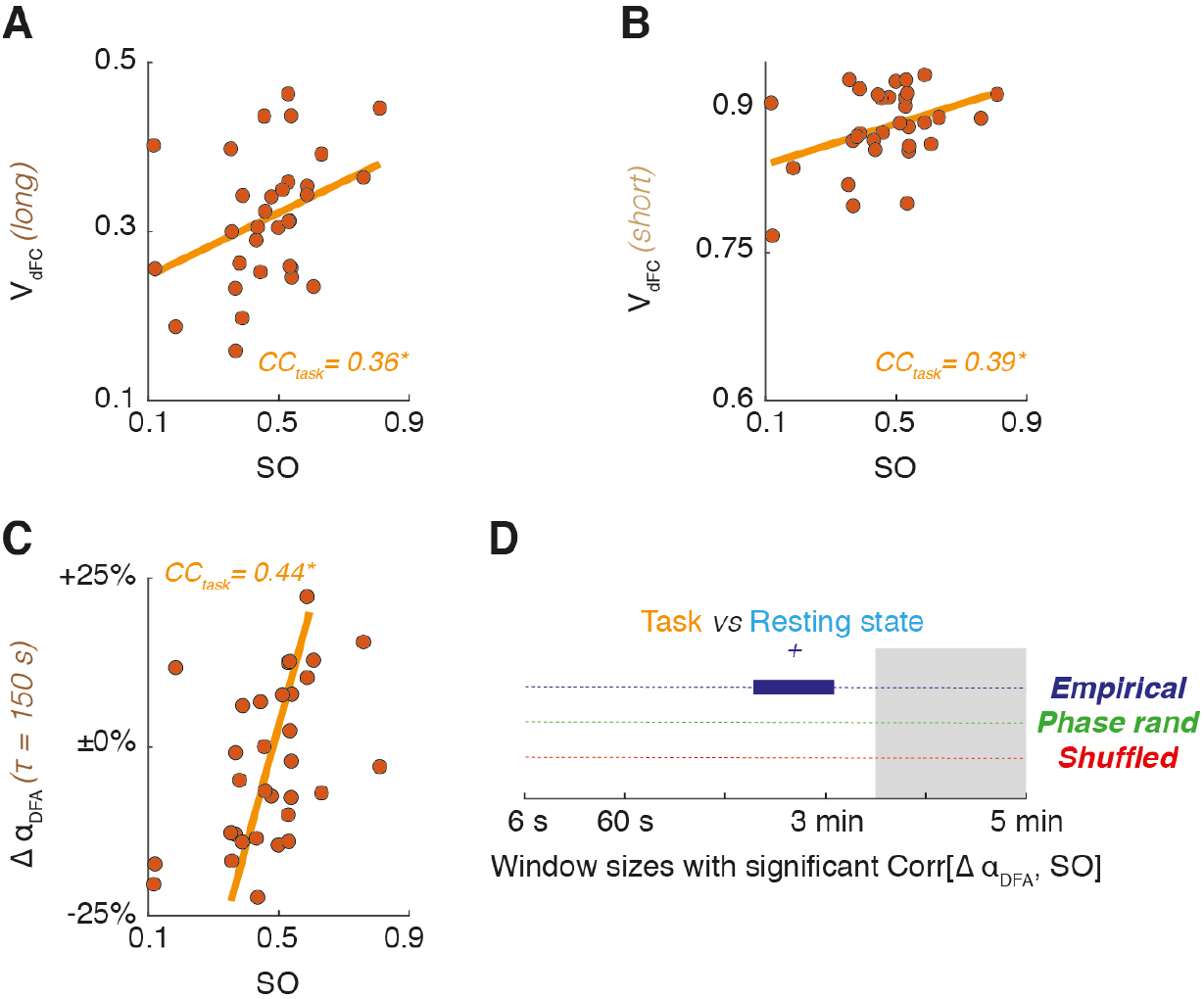
Correlation between dFC and cognitive/visuo-motor performance *(additional analyses)*. (A-B) The dFC speed of task dFC streams positively correlates with performance in a bimanual visuomotor task, tracked by a Spectral Overlap (SO). Shown here are a scatter plot for dFC speeds pooled in the long window sizes range (A) and in the short window sizes range (B), confirming the result of Figure 6A for intermediate window sizes range. (C-D) The variation of α_DFA_ between resting-state and task blocks, Δα_DFA_ = α_DFA_^task^-α_DFA_^rest^, positively correlates with SO during task (bootstrap with replacement confidence intervals for Pearson correlation: *, *p* < 0.05), as shown by (C) the scatter plot for a representative single window size in the long window sizes range. Such positive correlation (“+” sign over the range) holds significantly (bootstrap, *p* < 0.05) within a selected range of long window sizes and only for empirical data but not for surrogate dFC streams.

### Supplementary tables

**Table S1.** Used cortical parcellation and abbreviations.

| Abbreviation | Cortical Area |
| --- | --- |
| ENT | Entorhinal cortex |
| PARH | Parahippocampal cortex |
| TP | Temporal pole |
| FP | Frontal pole |
| FUS | Fusiform gyrus |
| TT | Transverse temporal cortex |
| LOCC | Lateral occipital cortex |
| SP | Superior parietal cortex |
| IT | Inferior temporal cortex |
| IP | Inferior parietal cortex |
| SMAR | Supramarginal gyrus |
| BSTS | Bank of the superior temporal sulcus |
| MT | Middle temporal cortex |
| ST | Superior temporal cortex |
| PSTC | Postcentral gyrus |
| PREC | Precentral gyrus |
| CMF | Caudal middle frontal cortex |
| POPE | Pars opercularis |
| PTRI | Pars triangularis |
| RMF | Rostral middle frontal cortex |
| PORB | Pars orbitalis |
| LOF | Lateral orbitofrontal cortex |
| CAC | Caudal anterior cingulate cortex |
| RAC | Rostral anterior cingulate cortex |
| SF | Superior frontal cortex |
| MOF | Medial orbitofrontal cortex |
| LING | Lingual gyrus |
| PCAL | Pericalcarine cortex |
| CUN | Cuneus |
| PARC | Paracentral lobule |
| INS | Insula |

Parcellation based on Desikan et al. (2006).

### Supplementary files

#### Software toolbox Sx. MATLAB toolbox for dFC calculations

A suite of MATLAB functions and scripts for performing dFC analyses analogous to the ones performed in this study can be downloaded as Supporting File Sx or at the web address *(waiting for release, it can be provided on request to authors)* with associated help and user guide.

## Notes

### Competing Interest Statement

The authors have declared no competing interest.

### Summary of Updates

With respect to v3, comparison with a control dataset ad a few minor changes have been introduced to satisfy peer reviewers. The v3 versions replaced a previous preprint submitted in 2017. All analyses had to be repeated following discovery of problems in preprocessing. All results were confirmed, but, meanwhile, our understanding of the methods had considerably progressed, leading to major changes in our presentation (also taking in account new analyses). This manuscript is thus the first of a fully new diptych containing a comprehensive presentation of our current vision of dFC as a random walk in FC space with complex spatiotemporal structure.

